# An AhR-Ovol1-Id1 regulatory axis in keratinocytes promotes skin homeostasis against atopic dermatitis

**DOI:** 10.1101/2024.01.29.577821

**Authors:** Zeyu Chen, Morgan Dragan, Peng Sun, Daniel Haensel, Remy Vu, Lian Cui, Yuling Shi, Xing Dai

**Affiliations:** Department of Dermatology, Shanghai Skin Disease Hospital, Tongji University School of Medicine, China; Department of Biological Chemistry, School of Medicine, University of California, Irvine, CA 92697, USA; The NSF-Simons Center for Multiscale Cell Fate Research, University of California, Irvine, CA 92697, USA; Department of Dermatology, Shanghai Tenth People’s Hospital, Tongji University School of Medicine, Shanghai, China; Institute of Psoriasis, Tongji University School of Medicine, Shanghai, China; Department of Dermatology, School of Medicine, University of California, Irvine, CA 92697, USA

**Keywords:** AhR, Ovol1, Id1, transcription factor, atopic dermatitis, epidermis, skin inflammation, skin barrier, IL-1, gamma delta T

## Abstract

Skin is our outer permeability and immune defense barrier against myriad external assaults. Aryl hydrocarbon receptor (AhR) senses environmental factors and regulates barrier robustness and immune homeostasis. AhR agonist is in clinical trial for atopic dermatitis (AD) treatment, but the underlying mechanism of action remains ill-defined. Here we report *OVOL1/Ovol1* as a conserved and direct transcriptional target of AhR in epidermal keratinocytes. We show that OVOL1/Ovol1 impacts AhR regulation of keratinocyte gene expression, and *Ovol1* deletion in keratinocytes hampers AhR’s barrier promotion function and worsens AD-like inflammation. Mechanistically, we identify Ovol1’s direct downstream targets genome-wide, and provide *in vivo* evidence for Id1’s critical role in barrier maintenance and disease suppression. Furthermore, our findings reveal an IL-1/dermal γδT cell axis exacerbating both type 2 and type 3 immune responses downstream of barrier perturbation in *Ovol1*-deficient AD skin. Finally, we present data suggesting the clinical relevance of OVOL1 and ID1 function in human AD. Our study highlights a keratinocyte-intrinsic AhR-Ovol1-Id1 regulatory axis that promotes both epidermal and immune homeostasis against AD-like inflammation, implicating new therapeutic targets for AD.

## INTRODUCTION

Barrier tissues such as skin reside at the interface of our body and the outside environment and protect us from a myriad of environmental aggressors such as pathogens and allergens. Tightly balanced proliferation and terminal differentiation of skin epidermal cells produce as well as maintain a physical permeability barrier, whereas skin resident (e.g., gamma delta T or γδT cells, dendritic cells or DCs) and infiltrating (e.g., neutrophils) immune cells provide an immunological defense barrier (*1, 2*). Despite frequent challenges from various external stimuli, skin maintains homeostasis under most conditions, owing to mechanisms that help resolve encounters without causing excessive tissue damage or inflammation. The regulatory mechanisms coordinating epidermal barrier maintenance with inflammatory response modulation at the epidermal-immune interface remain an active yet under-explored area of investigation.

Atopic dermatitis (AD) is the most common inflammatory skin disease that affects 15-20% of children and up to 10% of adults (*3*). Defective epidermal barrier caused by genetic mutations (e.g., *FLG*) and/or inflammatory cytokines is a salient feature, whereas environmental allergens such as house dust mites (HDM) are common triggers, of AD (*4–9*). Over-activated immune responses play a critical role in AD pathogenesis (*5*). Type 2 immunity, which is characterized by the production of T helper 2 (Th2)-related cytokines (e.g., IL-4, IL-13) and activation of eosinophil/mast cells (*10*), contributes to AD by impairing the epidermal barrier and causing symptoms such as inflammation and itch (*7–9, 11–13*). Emerging evidence suggests that type 3 immunity, characterized by the production of Th17-related cytokines (e.g., IL-17A) and recruitment of neutrophils, is also involved in AD development, especially in intrinsic, pediatric or Asian patients, by fueling the type 2 immune response (*5, 10, 14, 15*). Compared with healthy skin, AD skin is usually heavily colonized by pathogenic bacteria *Staphylococcus aureus*, which further aggravate epidermal barrier and immune defects to perpetuate disease progression (*4*). Biologics targeting IL-4/IL-13 or small molecule inhibitors targeting Janus kinase (JAK)/signal transducers and activators of transcription (STAT) signaling have shown great efficacy in treating AD patients (*16, 17*), but not all patients respond well to them. Development of additional therapies necessitates a better understanding of the molecular and cellular control of AD-associated barrier dysregulation and inflammation. In particular, epidermal-intrinsic mechanisms that enhance barrier strength while regulating the early innate immunological responses to AD-associated allergen/pathogen attacks are ideal therapeutic targets, but remain largely elusive.

Aryl hydrocarbon receptor (AhR), an environment-sensing xenobiotic receptor and ligand-activated transcription factor vital for epidermal barrier maintenance and skin immune homeostasis has attracted attention as a therapeutic target against both AD and psoriasis (*18–22*). However, based on animal model studies and patient clinical trials, the functional outcomes of AhR activation in AD-like skin inflammation appear complex and context (e.g., ligand)-dependent (*23–25*). Moreover, the cellular and molecular mechanisms underlying AhR function remain to be fully understood. Chemical activation of AhR in cultured normal human epidermal keratinocytes (NHEKs) upregulates *OVOL1*, a zinc finger transcriptional repressor gene identified by genome-wide association studies to be an AD and acne risk locus (*26–28*). *OVOL1* promotes the expression of epidermal differentiation-associated proteins such as filaggrin and loricrin in cultured NHEKs (*26*), whereas mouse *Ovol1* promotes epidermal cell cycle arrest and suppresses psoriasis-like skin inflammation (*29–32*). Whether *OVOL1/Ovol1* is directly activated by AhR and in turn regulates AD-like skin inflammation *in vivo* has not been investigated.

In this study, we identify *OVOL1*/*Ovol1* as a direct transcriptional target of AhR that in turn modifies AhR regulation of keratinocyte gene expression and epidermal barrier function. We show that keratinocyte-specific deficiency of *Ovol1* exacerbates AD-like inflammation induced by treatment with HDM and *Staphylococcus aureus*-derived toxin, staphylococcal enterotoxin B (SEB), and that Ovol1 does so by repressing the expression of myriad downstream targets but particularly transcription factor-encoding *Id1*. Using chemical inhibition and antibody blocking approaches, we uncover differential roles of Id1 and IL-1 signaling in *Ovol1* deficiency-associated barrier integrity and in orchestrating the downstream immune responses that involve dermal γδT cells and neutrophils. Finally, we present data to support the clinical relevance of OVOL1 and ID1 function in human AD. Our study underscores a keratinocyte-intrinsic AhR-Ovol1-Id1 regulatory axis that functions at the environment-barrier-immune interfaces to protect skin homeostasis against AD-like inflammation, as well as implicates new therapeutic targets for human AD.

## RESULTS

### AhR directly activates *OVOL1* expression in human keratinocytes to suppress cell cycle and promote differentiation

To explore the mechanism underlying the responsiveness of *OVOL1* expression to AhR activation, we examined the sequence of *OVOL1* promoter and identified several AhR-binding motifs (GCGTG) (Fig. 1A). ChIP-qPCR analysis in NHEKs showed that indeed AhR physically binds to these sites (Fig. 1B). Binding to *CYP1A1*, a well-known AhR downstream target (*33*), was also observed (Fig. 1B). Furthermore, small interfering RNA (siRNA)-mediated silencing of *AHR* expression in NHEKs resulted in significant downregulation of not only *AHR* canonical targets *CYP1A1* and *CYP1B1* (*33*), but also *OVOL1,* mRNA expression (Fig. 1C). These results demonstrate that *OVOL1* is a direct target of endogenous AhR protein in human keratinocytes.

**Figure 1.**
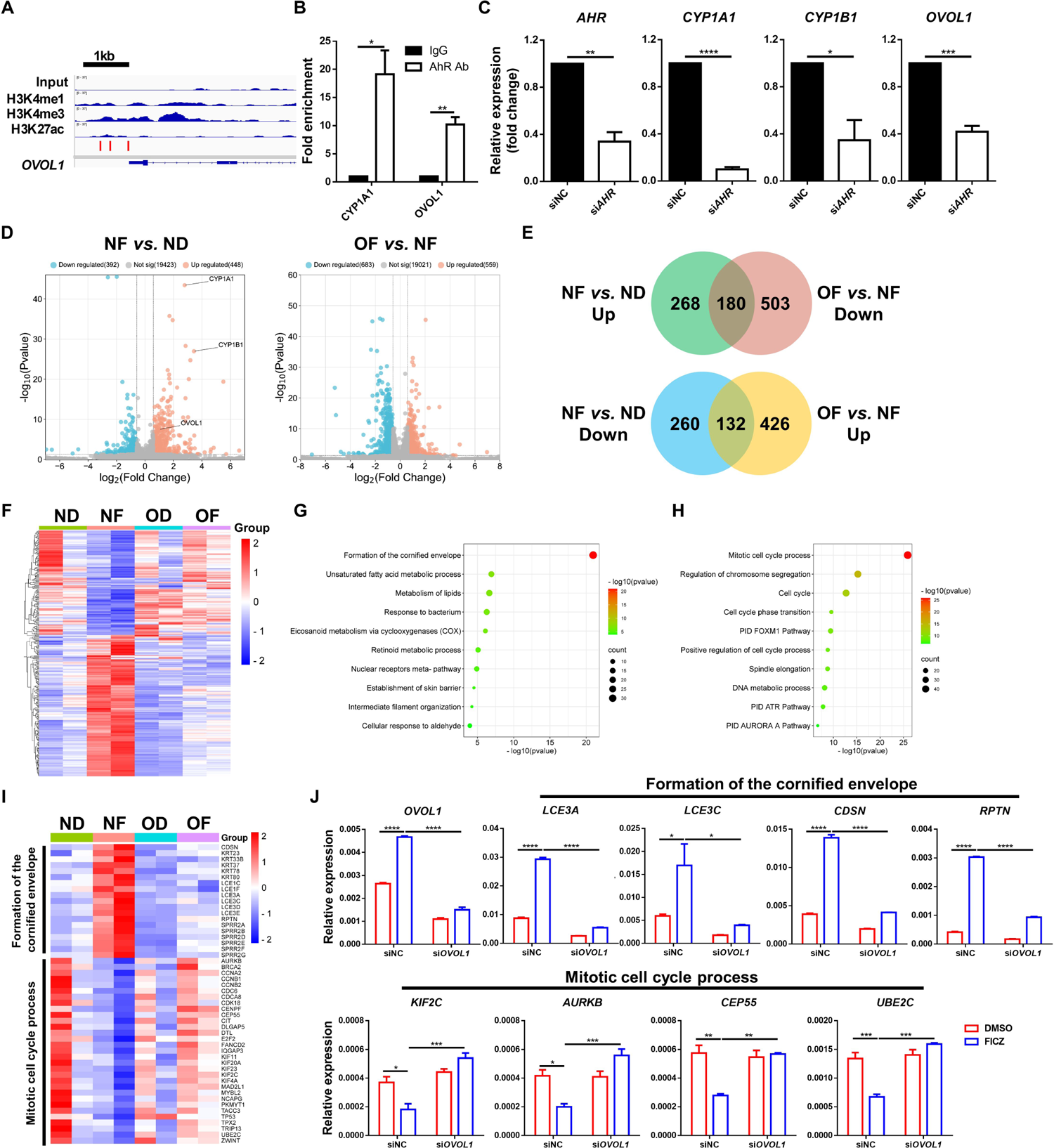
*OVOL1* is a target of AhR and modifies its regulation of gene expression in human keratinocytes. **(A)** Genome browser track for the indicated ChIP-seq signals across the *OVOL1* locus. Red bar indicates the presence of AhR binding motif (GCGTG). **(B)** ChIP-qPCR results of the indicated genes in FICZ-treated NHEKs. IgG control values were normalized to 1. Results are summarized from 3 independent experiments. **(C)** RT-qPCR results of the indicated genes in scrambled or *AHR* siRNA-treated NHEKs. Results are summarized from 3 independent experiments. **(D)** Volcano plots showing differential gene expression in the indicated samples. ND: siNC (negative control) + DMSO; NF: siNC + FICZ; OD: si*OVOL1* + DMSO; OF: si*OVOL1* + FICZ. Results are summarized from 2 independent experiments. **(E)** Venn diagrams of differentially expressed genes (DEGs) showing overlap between FICZ-induced genes (NF *vs.* ND) and OVOL1-dependent genes (OF *vs.* NF). UP, upregulated. Down, downregulated. Numbers of DEGs are as indicated. **(F)** Heatmap of the DEGs from (E). **(G-H)** Ingenuity pathways analysis of the 180 (G) or 132 (H) overlapping DEGs from (E). **(I)** Heatmap of DEGs in the indicated pathways. **(J)** RT-qPCR results of the indicated genes in DMSO- or FICZ-treated NHEKs with or without *OVOL1* knockdown. Results are summarized from 3 independent experiments. For (B), (C) and (J), data are mean + SEM. * *p* < 0.05, ** *p* < 0.01, *** *p* < 0.001, **** *p* < 0.0001. *p* values were calculated using 2-tailed unpaired Student *t* test (B and C) or two-way ANOVA (J).

Next, we assessed the importance of OVOL1 in mediating AhR function in human keratinocytes using an siRNA knockdown approach. Treatment of NHEKs with AhR agonist 6-formylindolo[3,2-b]carbazole (FICZ) led to an increased and decreased expression of 448 and 392 genes, respectively, in the presence of a scrambled siRNA (Fig. 1D, S1A, Table S1). As expected, *OVOL1*, *CYP1A1*, and *CYP1B1* were among the elevated genes. Metascape analysis of the FICZ-induced genes revealed “skin development”, “skin epidermis development”, and “metabolism of lipids” as top terms (Fig. S1B, Table S2), while the FICZ-suppressed genes were enriched for “mitotic cell cycle process”, “cell cycle”, and “positive regulation of cell migration” terms (Fig. S1C, Table S3). Knockdown of *OVOL1* in DMSO-treated NHEKs increased the expression of 304 genes and decreased the expression of 542 genes (Fig. S1D-E, Table S4). The downregulated genes were enriched for “skin development”, “positive regulation of hydrolase activity”, and “metabolism of lipids” (Fig. S1F, Table S5), while the upregulated genes were enriched for “cytokine signaling in immune system”, “network map of SARS-CoV-2 signaling pathway”, and “rhythmic process” (Fig. S1G, Table S6). These results reveal both overlapping (skin development and lipid metabolism) and divergent (cell cycle control *vs.* immune modulation) functions of AhR and OVOL1 in human keratinocytes.

We then examined how FICZ shapes OVOL1 function and vice versa. We found knockdown of *OVOL1* in FICZ-treated NHEKs to increase the expression of 559 genes and decrease the expression of 683 genes (Fig. 1D, S1E, Table S7). The downregulated genes were enriched for “keratinization”, “NOD-like receptor signaling pathway”, and “IL-18 signaling pathway” (Table S8). The upregulated genes were enriched for “mitotic cell cycle”, “cell cycle”, and “positive regulation of cell cycle process” (Table S9). These results suggest a partial shift in the molecular function of OVOL1, from predominantly regulating immune activity under homeostatic conditions to suppressing cell cycle when AhR signaling is activated by an exogenous ligand. Of the 448 genes upregulated and 392 genes downregulated by FICZ alone, 40.2% (180/448) and 33.7% (132/392), respectively, were dependent on *OVOL1* expression (Fig. 1E-F, Table S10). Interestingly, FICZ induction of *CYP1A1*, but not *CYP1B1*, was dependent on OVOL1 (Table S10). The top enriched pathways in the FICZ-induced, *OVOL1*-dependent genes were “formation of the cornified envelop”, “unsaturated fatty acid metabolic process”, and “metabolism of lipids” (Fig. 1G, I, Table S11). The top enriched terms in the FICZ-suppressed, *OVOL1*-dependent genes were “mitotic cell cycle process”, “regulation of chromosome segregation”, and “cell cycle” (Fig. 1H, I, Table S12). RT-qPCR analysis confirmed the failure of FICZ to upregulate genes in the “formation of the cornified envelop” pathway (e.g., *LCE3A*, *LCE3C*, *CDSN* and *RPTN*) and to downregulate genes in the “mitotic cell cycle process” pathway (e.g., *KIF2C*, *AURKB*, *CEP55* and *UBE2C*) when *OVOL1* is depleted from the keratinocytes (Fig. 1I-J). These data show that activated AhR signaling in human keratinocytes requires OVOL1 to promote the expression of genes associated with epidermal differentiation and lipid metabolism and to inhibit the expression of genes associated with cell cycle.

Collectively, our findings suggest a mechanistic model where AhR directly activates the expression of *OVOL1*, which in turn mediates AhR’s function in reprograming keratinocyte gene expression from pro-proliferation to pro-differentiation.

### AhR activates *Ovol1* expression in mouse keratinocytes, and requires *Ovol1* for downstream function *in vivo* in promoting barrier robustness against AD stimuli

The dramatic structural differences between human and mouse skin (*34, 35*) suggest that conservation of regulatory mechanisms cannot be assumed. This is a particularly important issue for AhR signaling given that observations based on mouse models are often used to infer clinical significance. To probe whether *Ovol1* is a direct target of AhR in mice, we analyzed the published AhR ChIP-seq data from primary murine keratinocytes (*36*), and found AhR to directly bind to the mouse *Ovol1* promoter (Fig. 2A). Furthermore, interrogation of published RNA sequencing (RNA-seq) data on primary keratinocytes from *Ahr*^+/+^ and *Ahr*^-/-^ mice (*37*) revealed significantly decreased expression of *Cyp1b1* and *Ovol1*, but not other known AhR targets such as *Cyp1a1, Bax* (*38*)*, Cdkn1b* (*39*)*, Gsta1, Gsta2* (*40*) *and Hsp90aa1* (*19*), in *Ahr*^-/-^ keratinocytes (Fig. 2B, S2A). Therefore, AhR direct activation of *Cyp1b1*/*CYP1B1* and *Ovol1/OVOL1*, though not all targets, in epidermal keratinocytes is likely evolutionarily conserved between mice and humans.

**Figure 2.**
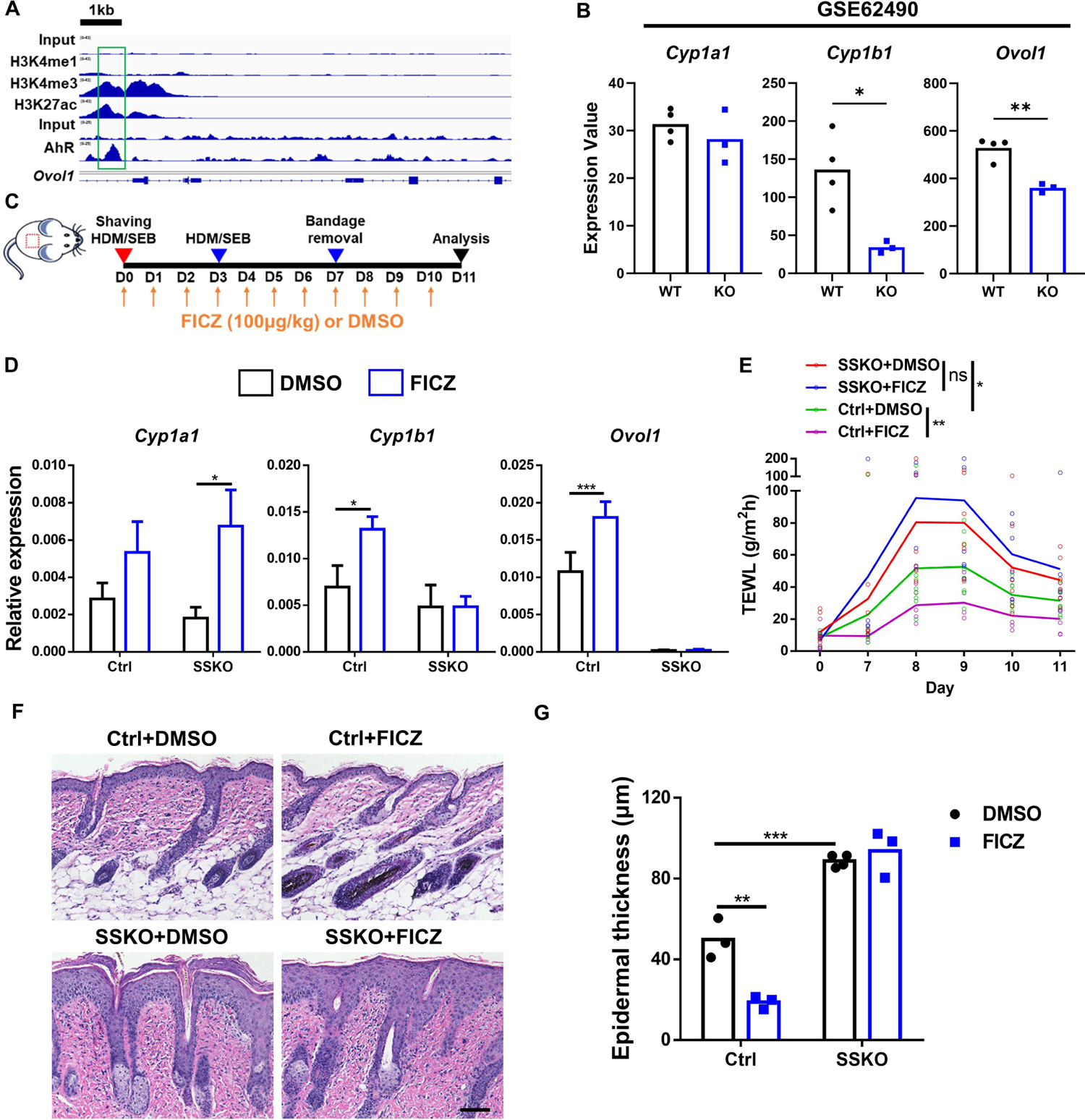
AhR requires *Ovol1* to mediate barrier maintenance in mice. **(A)** Genome browser tracks for the indicated ChIP-seq signals across the *Ovol1* locus. Green box highlights the AhR-bound region. **(B)** RNA-seq results of the indicated genes in mouse primary keratinocytes from *Ahr*^+/+^ (n = 4) or *Ahr*^-/-^ (n = 3) mice. **(C)** Experimental design for FICZ treatment in (D-G). DMSO was used as a vehicle control. **(D)** RT-qPCR results of the indicated genes in the whole skin at day 11. Data are mean + SEM. n = 4 mice per group. **(E)** Time course showing TEWL measurements. n = 5-9 mice per group. **(F)** Representative skin histology (H/E staining) at day 11. Scale bar = 100 μm. **(G)** Quantification of epidermal thickness. n = 3-4 mice per group. For (D), data are mean + SEM. * *p* < 0.05, ** *p* < 0.01, *** *p* < 0.001. ns, non-significant. *p* values were calculated using 2-tailed unpaired Student *t* test (B) or two-way ANOVA (D, E and G).

Next, we asked whether *Ovol1* is required for AhR activation of downstream target genes *in vivo*. We administered FICZ or DMSO via intraperitoneal (i.p.) injection to skin-epithelia-specific *Ovol1* knockout (SSKO: *K14*-Cre;*Ovol1*^f/-^) mice as well as control littermates (Fig. 2C). FICZ significantly elevated *Cyp1b1* expression in skin of control mice (Fig. 2D), indicating effectiveness of the treatment. *Ovol1* expression was also significantly upregulated by FICZ (Fig. 2D). Importantly, epidermal deletion of *Ovol1* (SSKO mice) abolished the FICZ-induced upregulation of *Cyp1b1* expression (Fig. 2D). However, FICZ did not significantly induce the expression of *Cyp1a1* in control littermate skin, while induction reached statistical significance in SSKO skin (Fig. 2D). Moreover, other known AhR targets including *Bax* (*38*)*, Cdkn1b* (*39*)*, Gsta1*, *Gsta2* (*40*) and *Hsp90aa1* (*19*), were not significantly induced in either control or SSKO skin (Fig. S2B). These data further highlight *Cyp1b1* and *Ovol1* as bona fide *in vivo* AhR targets in mouse skin, whereas AhR regulation of the other previously identified target genes is likely less robust or more complex. Although defining the precise mechanism underlying how Ovol1 contributes to AhR induction of *Cyp1b1* lies outside the scope of the current study, our ChIP-qPCR and ChIP-seq (see below) experiments revealed strong Ovol1 binding to the *Cyp1b1* promoter (Fig. S2C-D), suggesting feedforward regulation (Fig. S2E).

AhR signaling in keratinocytes promotes barrier integrity against AD-associated perturbations (*19, 20, 23, 41*), while AhR’s expression is significantly decreased in AD epidermis (Fig. S2F) per interrogation of published RNA-seq data (*42*). To assess whether AhR’s barrier-promoting function is dependent on *Ovol1* expression in keratinocytes, we turned to an AD mouse model induced by HDM and SEB, two agents that are clinically related to human AD (*4*) and known to induce AD-like phenotypes in mice when used in combination (*43*). As expected, applying FICZ to HDM/SEB-treated mice partially normalized their trans-epidermal water loss (TEWL) in control mice (Fig. 2E). However, FICZ failed to rescue the exacerbated barrier dysfunction in SSKO mice (Fig. 2E; we will return to the effect of *Ovol1* deletion itself in the next section). Histological analysis of skin sections showed that FICZ treatment significantly reduced epidermal thickness in HDM/SEB-treated control mice but not in SSKO mice (Fig. 2F-G). Collectively, these data provide the first *in vivo* evidence that AhR-mediated promotion of barrier function and suppression of epidermal proliferation in skin treated with AD stimuli require an intact *Ovol1* gene in epidermal keratinocytes.

### *Ovol1* in keratinocytes protects skin from AD-like barrier dysregulation and pathology

Our discovery of *Ovol1* as a direct and functionally significant downstream target of AhR in keratinocytes raises an important question of whether *Ovol1* plays a regulatory role in the development of AD-like skin inflammation, and if so, what the underlying mechanisms is. Towards addressing this, we analyzed published single-cell RNA-seq data of whole skin samples from healthy individuals and AD patients (*44*), and found downregulated *OVOL1* expression in AD keratinocytes, especially in lesional skin (Fig. 3A, S3A). In situ detection of *OVOL1* RNA validated its reduced expression in AD lesional skin epidermis (Fig. 3B). Supporting the functional relevance of this observation, deletion of *Ovol1* from mouse keratinocytes significantly aggravated the HDM/SEB-induced epidermal phenotypes, evident through the more severe barrier disruption and epidermal hyperplasia in SSKO mice compared with control littermates (Fig. 2E-G). These findings held true when the experiments were repeated in the absence of DMSO treatment (Fig. 3C-H). Not only the TEWL values were higher and epidermal thickening more prominent in HDM/SEB-treated SSKO mice, but also the other AD-like skin phenotypes, as measured by clinical score of skin eruption, scaling, bleeding and redness as well as dermal thickening, were significantly more severe in SSKO mice than in control littermates (Fig. 3C-H, S3B). At a histological level, lesional skin of SSKO mice, but not their control littermates, showed AD-like pathology including spongiosis and epidermal infiltration of eosinophils (Fig. 3G).

**Figure 3.**
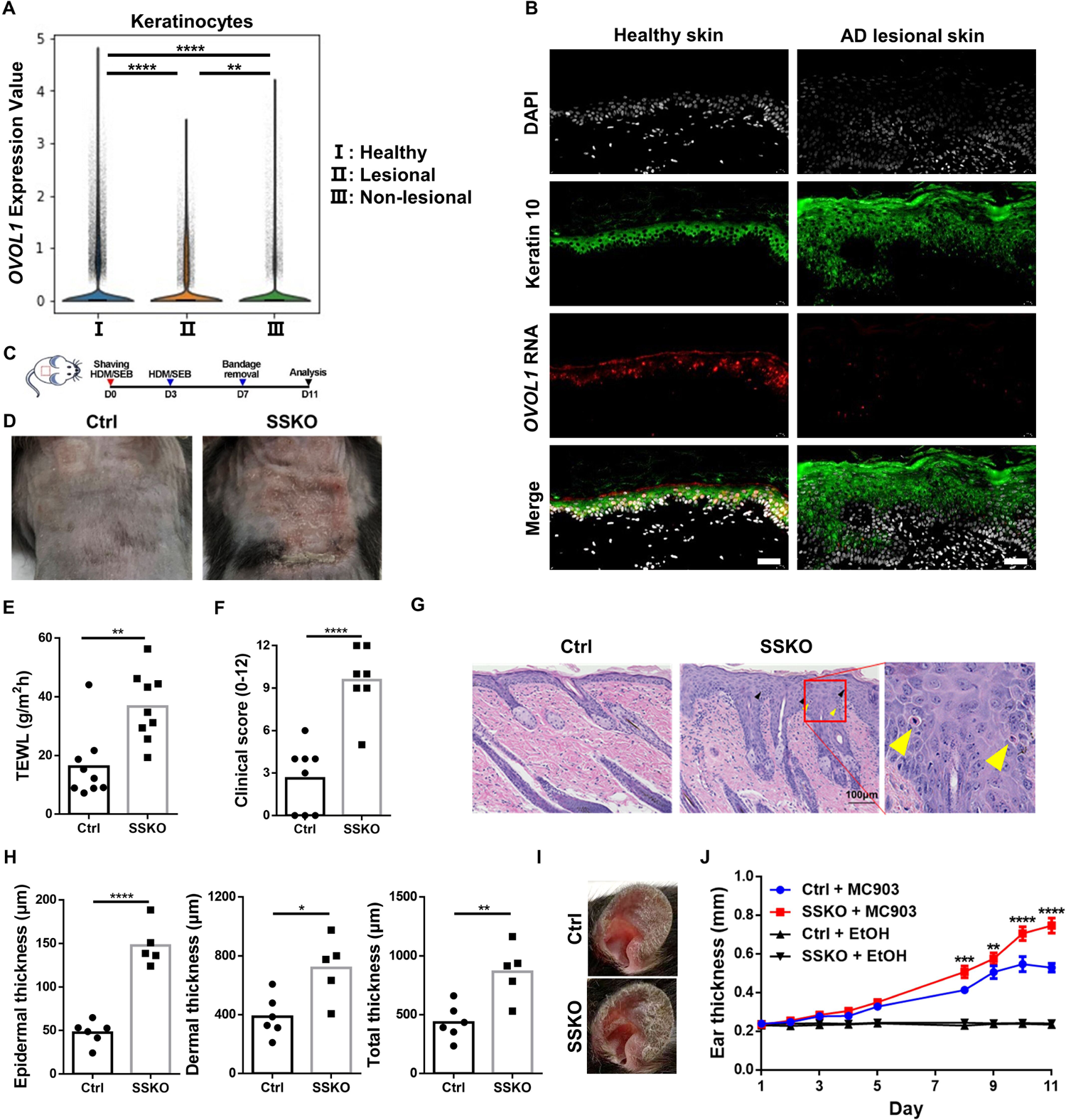
*Ovol1* deficiency in keratinocytes aggravates AD-like skin inflammation. **(A)** Violin plots of *OVOL1* expression in keratinocytes from healthy, lesional, and non-lesional skin. **(B)** Expression of keratin 10 protein and *OVOL1* RNA in healthy and lesional skin (n = 4 per group). DAPI stains the nuclei. Scale bar = 40 μm. **(C)** Experimental design for HDM/SEB treatment in (D-H). **(D)** Representative photographs of HDM/SEB-treated control (Ctrl) and SSKO mice at day 11. **(E)** Clinical scores of HDM/SEB-treated Ctrl (n = 8) and SSKO (n = 7) mice. **(F)** TEWL values of HDM/SEB-treated Ctrl (n = 9) and SSKO (n = 9) mice. **(G)** Representative skin histology (H/E staining) at day 11. Scale bar = 100 μm. Black arrowheads indicate spongiosis. Yellow arrowheads indicate epidermal infiltration of eosinophils. **(H)** Quantification of epidermal, dermal and total skin thickness in Ctrl (n = 6) and SSKO (n = 5) mice. **(I)** Representative photographs of MC903-treated Ctrl and SSKO mice at day 11. **(J)** Ear thickness (mean ± SEM) of MC903-treated Ctrl (n = 6) and SSKO (n = 6) mice. * *p* < 0.05, ** *p* < 0.01, *** *p* < 0.001, **** *p* < 0.0001. *p* values were calculated using *t* test followed by Benjamini-Hochberg correction (B) or 2-tailed unpaired Student *t* test (F, G and I) or two-way ANOVA (K, Ctrl + MC903 *versus* SSKO + MC903).

Because AD is a chronic skin disease, we next tested whether *Ovol1*’s protective role persists after prolonged exposure to HDM and SEB. After 3 rounds of HDM/SEB treatment, barrier function was no longer different between control and SSKO mice, but the skin clinical symptoms and increased epidermal/dermal thickness were still more remarkable in SSKO mice than control littermates (Fig. S3C-G). This result implicates a long-term protective role of *Ovol1* against AD-like skin symptoms beyond simply preserving a physical barrier.

We also examined the impact of keratinocyte-specific *Ovol1* deletion in another widely used AD model, namely ear skin responses to MC903, a vitamin D analogue that induces AD-like skin pathology (*45*). Compared to control littermates, the ear skin of MC903-treated SSKO mice was more symptomatic and significantly thicker (Fig. 3I-J). In a similar vein, the ears of *Ovol1*^-/-^ mice (*30, 31*) also showed worsened skin inflammation and were significantly thicker than that of control littermates (Fig. S3H-I). When a high dose of MC903 was used on the right ears, even the left ear skin (sham control) and back skin of *Ovol1*^-/-^ mice were thicker and more proliferative than in control littermates (Fig. S3J-L), suggesting that local MC903 administration may have elicited a systemic effect in these mice. Interestingly, the serum levels of Cxcl2 and G-CSF, factors involved in neutrophil recruitment or maturation, were significantly elevated in *Ovol1*^-/-^ mice compared to control littermates (Fig. S3M). Moreover, neutrophil presence was observed in skin of both the directly applied (right) and sham control (left) ears of MC903-treated *Ovol1*^-/-^ mice (Fig. S3N-O).

Taken together, our data demonstrate the functional importance of keratinocyte-expressed *Ovol1* in suppressing agent-induced AD-like skin pathology in two different experimental models, and they show that this protective effect of *Ovol1* takes effect under both acute and chronic perturbations, and against both local and systemic effects.

### Identification of novel downstream targets of Ovol1 in keratinocytes, and *Id1* as a functional target that mediates Ovol1’s barrier-associated function

To probe how Ovol1 protects against AD-like barrier dysregulation and skin pathology, we first sought to identify its direct targets using ChIP-seq in primary mouse keratinocytes (*32*). Using MACS2 broad peak calling, a total of 1,109 Ovol1-bound peaks were identified that passed a significance threshold (q value < 0.05) (Table S13). A large fraction (73%) of the peaks were found to reside in gene promoter regions within 1 kb of the transcription start site (TSS) (Fig. 4A-B), suggesting preferential binding of Ovol1 to proximal gene promoters. Substantial binding to distal intergenic sequences (10.6 %) was also seen (Fig. 4B). Homer analysis revealed that ∼50% of the Ovol1-bound loci contain at least one consensus binding motif (CCGTTA) (Fig. 4C), which is identical to the consensus sequence previously identified for recombinant Ovol1 binding using an in vitro site selection assay (*29*). Interestingly, 42.6% of the Ovol1-bound loci contain TTTTCGCG (Fig. 4C), a consensus motif for E2F transcription factor binding, suggesting that Ovol1 may also be recruited to a subset of its cognate promoters *in vivo* through binding to E2F sequences and/or through E2F/Rb complexes (*46*). Metascape analysis of the Ovol1-bound genes revealed “tube morphogenesis”, “regulation of cell projection organization”, “regulation of cytoskeleton organization”, “regulation of mRNA stability”, “positive regulation of organelle organization”, and “regulation of cell cycle process” as top terms (Fig. 4D; Table S14), suggesting a role of Ovol1 in directly regulating these cellular processes.

**Figure 4.**
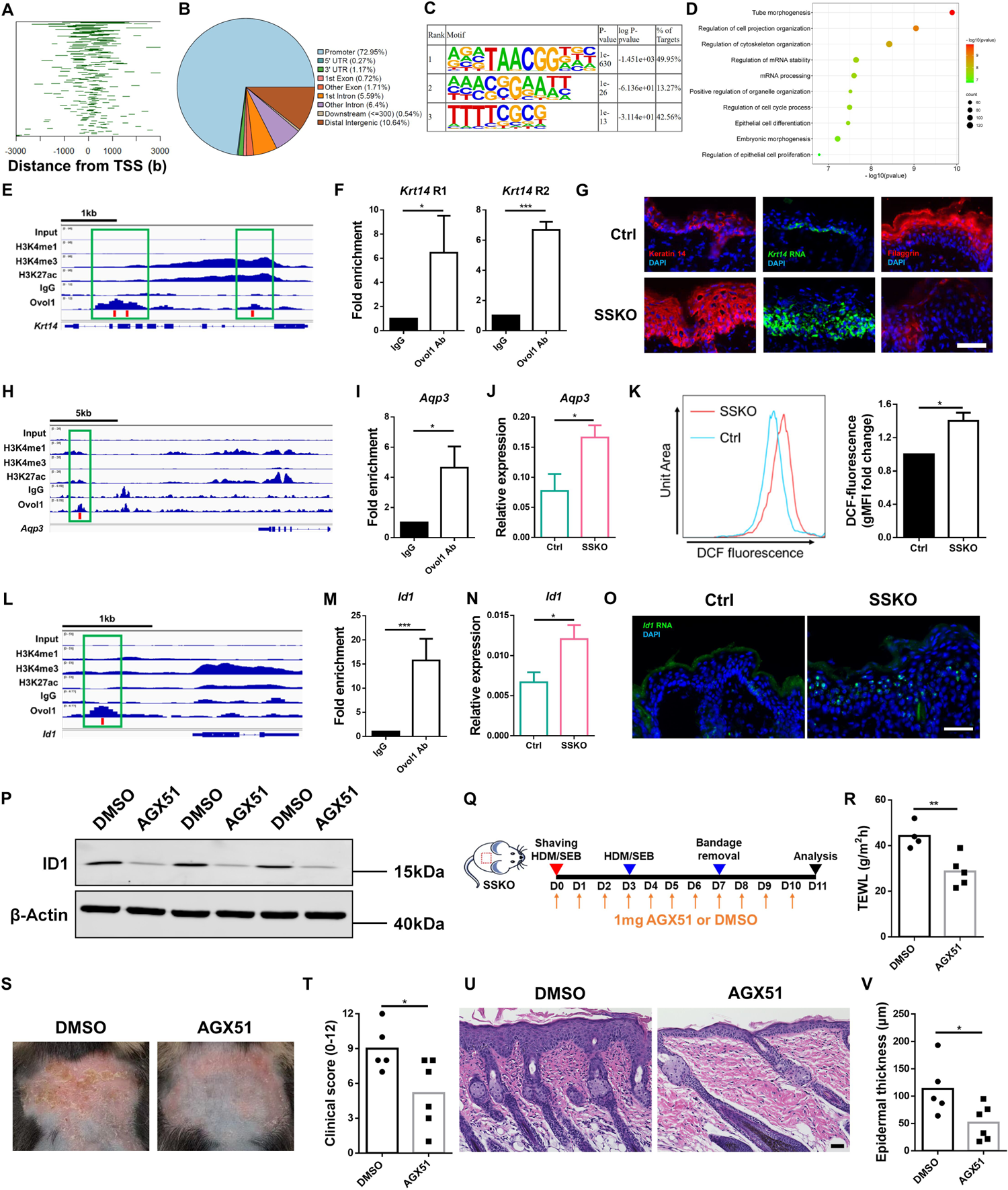
Identification of direct and functional targets of Ovol1. **(A)** Line plots of the distance between the Ovol1 ChIP-seq peaks and the TSS. **(B)** Pie charts depicting annotated genomic features of the called ChIP-seq peaks. **(C)** Homer motif analysis showing the predominance of the Ovol consensus motif in the ChIP-seq peaks. **(D)** Metascape analysis for all the genes associated with peaks for Ovol1. **(E)** Genome browser track for the indicated ChIP-seq signals across the *Krt14* locus. **(F)** ChIP-qPCR results of *Krt14* in mouse keratinocytes treated with Ca^2+^ (1.8 mM) for 24 hours. IgG control values were normalized to 1. Data are mean + SEM. Results are summarized from 3 independent experiments. **(G)** Representative immunofluorescence images of keratin 14 protein, filaggrin protein, and RNAScope data of *Krt14* mRNA in lesional skin of HDM/SEB-treated Ctrl and SSKO mice at day 11. n = 3 mice per group. Scale bar = 50 μm. **(H)** Genome browser track for the indicated ChIP-seq signals across the *Aqp3* locus. **(I)** ChIP-qPCR results of *Aqp3*. See (F) for details. **(J)** RT-qPCR result of *Aqp3* expression in whole skin of HDM/SEB-treated Ctrl (n = 6) and SSKO (n = 5) mice at day 11. Data are mean + SEM. **(K)** Geometric mean fluorescence intensity (gMFI) values of DCF fluorescence in epidermal keratinocytes from HDM/SEB-treated Ctrl and SSKO mice at day 11. Values of Ctrl mice were normalized to 1. n = 3 mice per group. **(L)** Genome browser track for the indicated ChIP-seq signals across the *Id1* locus. Red bars in (E), (H), and (L) indicate the presence of Ovol1 binding motif (CCGTTA), and green boxes highlight the Ovol1-bound regions. **(M)** ChIP-qPCR results for *Id1*. See (F) for details. **(N)** RT-qPCR result of *Id1*. See (J) for details. **(O)** Representative RNAScope data of *Id1* mRNA in lesional skin of HDM/SEB-treated Ctrl and SSKO mice at day 11. n = 3 mice per group. Scale bar = 50 μm. DAPI in (G) and (O) stains the nuclei. **(P)** Western blotting results of ID1 expression in HaCaT cells treated with DMSO or AGX51 (100 μM) for 24 hours. Results are summarized from 3 independent experiments. **(Q)** Experimental design for AGX51 treatment in (R-V). DMSO was used as a vehicle control. **(R)** TEWL values of DMSO- or AGX51-treated SSKO mice at day 11. n = 4-5 mice per group. **(S)** Representative photographs of DMSO- or AGX51-treated SSKO mice at day 11. **(T)** Clinical scores of DMSO- or AGX51-treated SSKO mice at day 11. n = 5-6 mice per group. **(U)** Representative skin histology (H/E staining) at day 11. Scale bar = 50 μm. **(V)** Quantification of epidermal thickness. n = 5-6 mice per group. Data are mean + SEM. * *p* < 0.05, ** *p* < 0.01, *** *p* < 0.001. *p* values were calculated using 2-tailed unpaired Student *t* test.

Delving deeper into Ovol1 regulation of the cytoskeleton, we noted direct binding of Ovol1 to several keratin-encoding genes including *Krt5*, *Krt14*, and *Krt10*. Keratin 14 is an epidermal basal (stem/progenitor) cell-specific keratin that not only supports basal cell structure but also regulates keratinocyte proliferation and differentiation (*47*). Our ChIP-seq detected Ovol1 binding at two regions in the *Krt14* locus, one of which lacks H3K27Ac, H3K4me1 and H3K4me3 histone modifications of active promoter/enhancers (72–75), where a total of three Ovol1-binding motifs reside (Fig. 4E). This binding was confirmed using ChIP-qPCR (Fig. 4F). Importantly, the expression of both *Krt14* mRNA and keratin 14 protein was considerably increased and expanded in the epidermis of HDM/SEB-treated SSKO relative to control mice (Fig. 4G). This increase, together with the dramatically reduced expression of terminal differentiation marker filaggrin in the suprabasal compartment of SSKO epidermis (Fig. 4G), suggests that *Ovol1*-deficient keratinocytes are arrested in a progenitor cell/early differentiation state at least in part due to loss of direct Ovol1 repression of *Krt14* expression.

To further identify functionally relevant Ovol1 targets, we cross-examined the Ovol1 ChIP-seq hits with genes that were upregulated upon *OVOL1* knockdown (Fig. S1D-E) or *Ovol1* deficiency. This comparison identified 17 Ovol1-bound genes (e.g., *Dmwd*, *Fosb*, *Ganab*, *Hic1*) whose human homologs showed increased expression in OVOL1-depleted NHEKs (Fig. S4A), and 39 Ovol1-bound genes (e.g., *Aqp3*, *Id1*, *Slpi*, *Stra6*) whose expression was increased in *Ovol1*-deficient mouse epidermis during the “innate immune response phase” of skin inflammation (*31*) (Fig. S4B). “Organic hydroxyl compound transport” (*Aqp3*, *Slc16a1* and *Stra6*) and “positive regulation of supramolecular fiber organization” (*Id1*, *Psrc1* and *Efemp2*) were terms enriched in the 39 mouse genes (Fig. S4C). Elevated expression of *AQP3* and *ID1* upon OVOL1 knockdown was observed in NHEKs with altered culturing condition (e.g., with tryptic soy broth) (Fig. S4D), leading us to zoom in on *Aqp3* and *Id1* as candidate Ovol1 targets in AD-like skin inflammation.

A putative enhancer ∼13 kb downstream of the *Aqp3* gene was found to contain an Ovol1-binding consensus, and ChIP-seq and ChIP-qPCR results showed that Ovol1 indeed binds to this site in mouse keratinocytes (Fig. 4H-I). RT-qPCR analysis detected significantly increased *Aqp3* expression in the lesional skin of HDM/SEB-treated SSKO mice relative to control skin (Fig. 4J). Since Aqp3 is known to promote H_2_O_2_ accumulation during skin inflammation (*48*), we compared H_2_O_2_ levels in keratinocytes of HDM/SEB-treated SSKO and control mice and found the level to be significant elevated in the former (Fig. 4K).

Transcription factor-encoding *Id1* is a known Ovol1 target in trophoblast cells, and has been shown to promote keratinocyte proliferation (*49–51*). Our ChIP-seq and ChIP-qPCR results revealed Ovol1 binding to a putative upstream enhancer of *Id1* in mouse keratinocytes, and this enhancer contains an Ovol1-binding consensus sequence (Fig. 4L-M). Moreover, RT-qPCR and RNAScope detected increased and expanded *Id1* expression in the lesional skin epidermis of HDM/SEB-treated SSKO mice compared to control counterparts (Fig. 4N-O). To determine whether aberrantly enhanced *Id1* expression functionally contributes to the disrupted skin barrier and pathology of SSKO mice, we utilized AGX51, a small chemical inhibitor that induces Id1 protein degradation *in vivo* and *in vitro* (*52*) (Fig. 4P). Compared to DMSO vehicle control, i.p administration of AGX51 significantly reduced TEWL and epidermal thickness, as well as improved the AD-like clinical score, of HDM/SEB-treated SSKO mice (Fig. 4Q-V, S4E-F). These results show that, despite Ovol1 directly binding and regulating myriad downstream targets, *Id1* is a functionally important target and its inhibition can partially rescue the epidermal barrier defect and associated pathology of AD-like skin in SSKO mice.

### Both type 2 and type 3 immune responses are enhanced in HDM/SEB-treated SSKO mice, and can be partially suppressed by functional inhibition of γδT cells

Although the importance of epidermal barrier in AD pathogenesis has been well-recognized, little is known about mechanistically how barrier dysregulation facilitates disease progression especially at the epidermal-innate immune interface (*14, 53*). To seek insights into the skin immunological responses to *Ovol1* deletion-instigated, AD-associated epidermal dysregulation, we used flow cytometry to compare the immune cell profiles in control and SSKO skin during homeostasis and AD-like inflammation. Langerhans cells (LCs) and dendritic epidermal T cells (DETCs) are immune cell types that reside within the mouse epidermis (*54*), and their relative abundance per total immune cells (CD45^+^) was not altered in the epidermis of SSKO mice during homeostasis (Fig. S5A-C). Upon stimulation with HDM and SEB, SSKO skin exhibited a trend of increase in the number of total immune cells relative to the control littermate skin (Fig. 5A-C). Specifically, SSKO skin contained significantly higher numbers of dermal γδT cells, neutrophils, and DCs than control skin, whereas the numbers of LCs, DETCs, macrophages, CD4^+^ T cells, CD8^+^ T cells, and innate lymphoid cells (ILCs) were not significantly affected (Fig. 5A-C, S5D-G). Of note, T cells (including dermal γδT cells) were the most abundant immune cell types in HDM/SEB-treated skin (Fig. 5A-C, S5F-G). In SSKO skin, there was also a significant increase in the number of CD11b^+^ cells not identifiable as LCs, macrophages, DCs, or neutrophils, and this population likely included eosinophils (Fig. 5A, 5C, S5F-G). Intriguingly, dermal γδT cells were still increased in SSKO mice treated with 3 rounds of HDM and SEB, while neutrophils and DCs were no longer significantly different from controls at this time (Fig. S5H-L).

**Figure 5.**
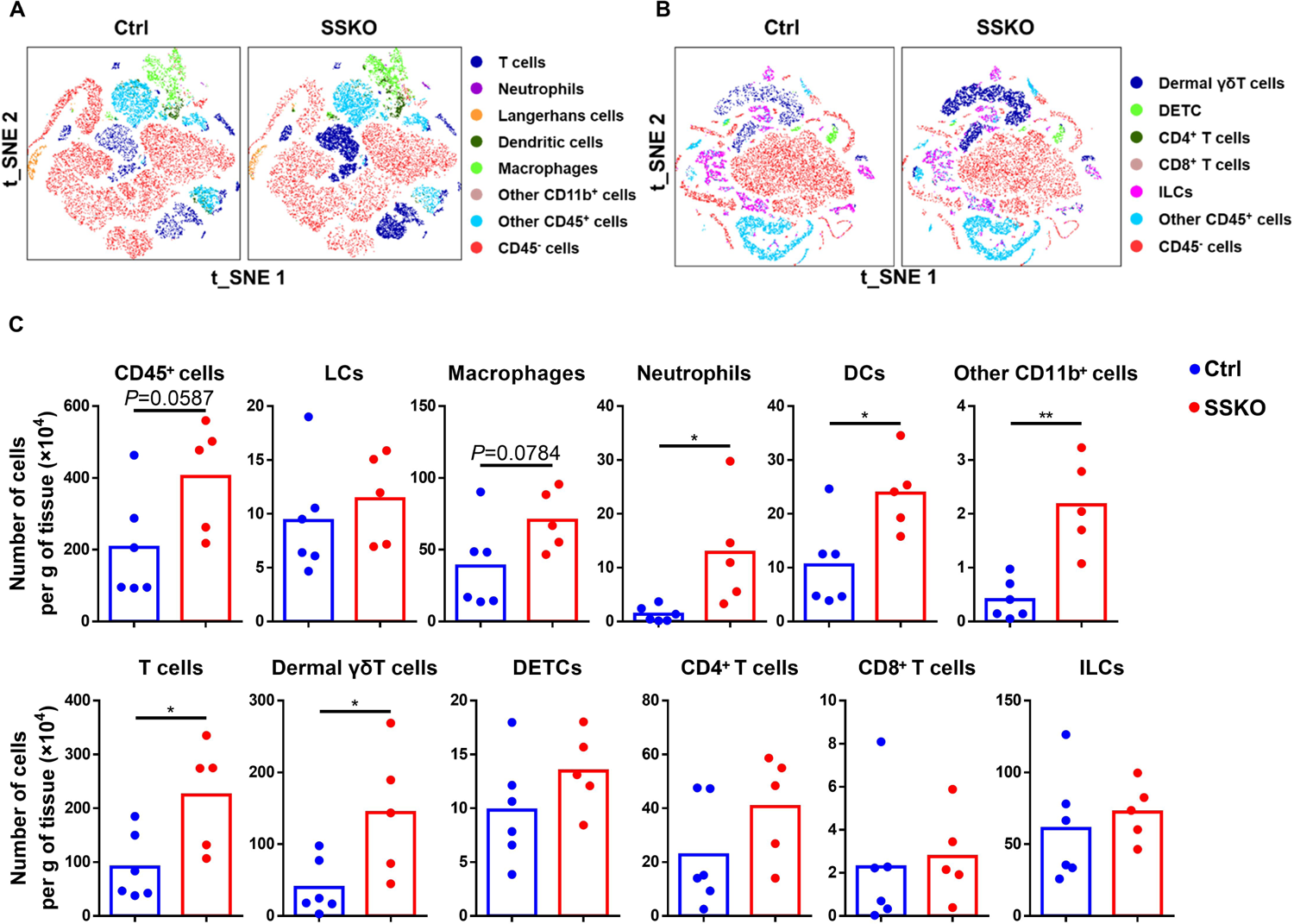
Immune cell profiles in HDM/SEB-treated control and SSKO skin. **(A-C)** Flow cytometry analysis of back skin of HDM/SEB-treated Ctrl and SSKO mice at day 11. Shown are tSNE visualization of the indicated cell types (A, B) and quantification of the number of the indicated immune cell types per gram of skin tissue (C). n = 6 for Ctrl; n = 5 for SSKO. * *p* < 0.05, ** *p* < 0.01, *** *p* < 0.001. *p* values were calculated using 2-tailed unpaired Student *t* test.

Dermal γδT cells express γδTCR and are an integral component of the skin local immunosurveillance program (*55*). They are kick-starters of inflammation and often the main producers of IL-17 in various models of inflammatory diseases especially in the early stages of inflammatory responses (*56, 57*). The dramatic and persistent increase of dermal γδT cells in barrier-deficient, AD-like SSKO skin led us to hypothesize that their accumulation contributes to the exacerbated inflammation and pathology. To test this, we administered γδTCR antibody to HDM/SEB-treated SSKO mice to functionally block γδT cells (*58–60*) (Fig. 6A). γδTCR antibody near-completely prevented flow cytometry detection of dermal γδT cells and DETCs (Fig. 6B), testifying the effectiveness of the treatment. Importantly, γδTCR antibody treatment significantly alleviated AD-like skin phenotypes in SSKO mice including clinical score, epidermal and dermal thickness (Fig. 6C-F, S6A-B), but did not significantly restore epidermal barrier function (Fig. 6G). Flow cytometry analysis revealed that the numbers of total immune cells, neutrophils, DCs, and T cells in AD-like SSKO skin were all significantly reduced by γδTCR antibody treatment (Fig. 6H, S6C-E). These data uncover a critical role of γδT cells (presumed to be dermal γδT cells because of their accumulation in SSKO skin) in orchestrating both innate (i.e., promoting neutrophil accumulation) and adaptive (i.e., promoting DC accumulation) immunity downstream of epidermal/barrier dysregulation in AD-like skin of SSKO mice.

**Figure 6.**
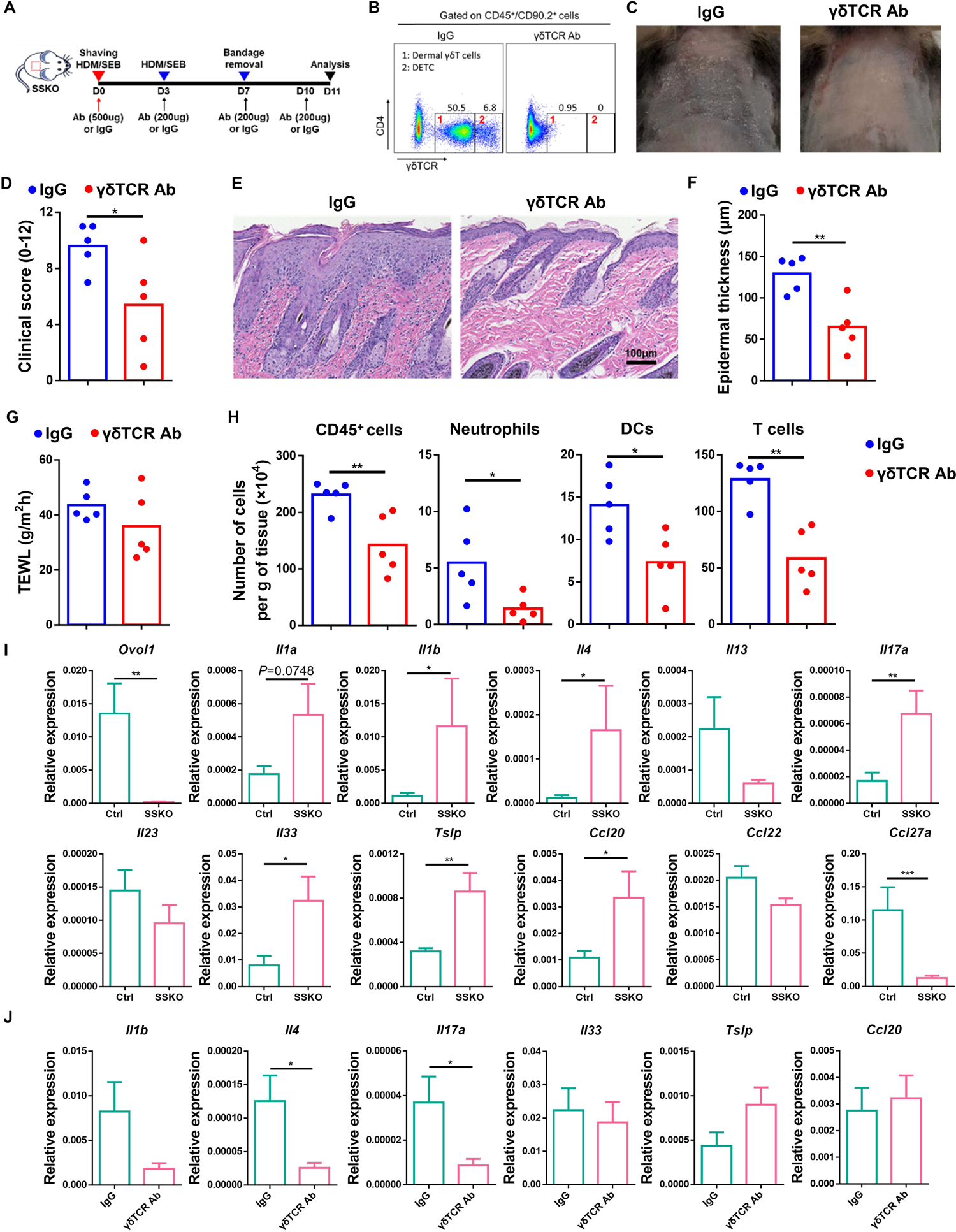
γδT cells contribute to the exacerbated skin inflammation in HDM/SEB-treated SSKO mice. **(A)** Experimental design for γδTCR antibody (Ab) treatment. IgG was used as a control. **(B)** Representative flow cytometry plots for dermal γδT cells (1) and DETCs (2) in CD45^+^CD90.2^+^ cells. Numbers in the plots represent relative abundance of the indicated cell populations. **(C)** Representative photographs of IgG- or γδTCR Ab-treated SSKO mice at day 11. **(D)** Clinical scores of IgG- or γδTCR Ab-treated SSKO mice at day 11. n = 5 mice per group. **(E)** Representative skin histology (H/E staining) at day 11. Scale bar = 100 μm. **(F)** Quantification of epidermal, dermal and total skin thickness. n = 5 mice per group. **(G)** TEWL values of IgG- or γδTCR Ab-treated SSKO mice at day 11. n = 5 mice per group. **(H)** Quantification of the number of the indicated cell types per gram of skin tissue. n = 5 mice per group. **(I, J)** RT-qPCR results of the indicated genes in whole skin of HDM/SEB-treated Ctrl (n = 6) and SSKO (n = 5) mice (I), or of IgG- or γδTCR Ab-treated SSKO mice (n = 5 mice per group) at day 11 (J). Data are mean + SEM. * *p* < 0.05, ** *p* < 0.01, *** *p* < 0.001. *p* values were calculated using 2-tailed unpaired Student *t* test or Mann-Whitney U test.

To elucidate the molecular changes underlying the enhanced AD-like skin phenotypes of SSKO mice, we examined the mRNA expression of type 2 and type 3 cytokines and chemokines in the HDM/SEB-treated lesional skin. As expected, *Ovol1* expression was abolished in SSKO skin (Fig. 6I). Importantly, the expression of *Il4*, *Il17a*, *Il33*, *Tslp* and *Ccl20* was significantly upregulated in SSKO skin, whereas the expression of *Il13*, *Il23, Ccl22* and *Ccl27a* was either unaffected or downregulated (Fig. 6I). These molecular profiles suggest that some, though not all, aspects of type 2 (*Il4, Il33* and *Tslp*), and type 3 (*Il17a* and *Ccl20*) immunity were enhanced in the AD-like skin by epidermal loss of *Ovol1*. Furthermore, RT-qPCR analysis revealed significant decreases in the expression of both *Il4* and *Il17a* in the AD-like skin lesions of γδTCR antibody-treated SSKO mice (Fig. 6J). Considering that dermal γδT cells are an appreciable source of IL-17 expression (*61*), we surmise that they also directly or indirectly contribute to the production of type 2 cytokine IL-4 and the potentiation of type 2 (Th2) immune responses in AD-like skin.

### Differential roles of IL-1 signaling and Id1 in regulating the epidermal barrier and immune response in HDM/SEB-treated SSKO skin

Next, we investigated the potential mechanism by which epidermal dysregulation in SSKO mice elicits the dermal γδT response. Epidermal barrier disruption can result in the elevated expression of alarmins such as IL-1α (*31, 62, 63*), and IL-1 signaling is known to activate dermal γδT cells during psoriasis-like skin inflammation in mice (*64, 65*). Indeed, we found the expression of *Il1a* to be near-significantly, and *Il1b* to be significantly, upregulated in SSKO lesional skin (Fig. 6I). Furthermore, we detected a significant increase of *Il1a* expression in the epidermis of HDM/SEB-treated SSKO mice compared to control littermates (Fig. 7A). Therefore, we asked whether enhanced IL-1 signaling might mediate the accumulation of excessive dermal γδT cells in the AD-like skin of SSKO mice. Specifically, we i.p administered an IL-1R antibody to HDM/SEB-treated SSKO mice to block IL-1 signaling (Fig. 7B). Flow cytometry analysis revealed that IL-1R antibody treatment led to a ∼2.5-fold decrease in the number of dermal γδT cells in skin of HDM/SEB-treated SSKO mice compared to IgG control-treated SSKO littermates (Fig. 7C-D, S7A-B). Consistent with this change, AD-like skin defects in HDM/SEB-treated SSKO mice were dramatically alleviated by the IL-1R antibody, presenting improved clinical score and reduced epidermal thickness (Fig. 7E-H, S7C). This said, TEWL and dermal thickness were not significantly affected (Fig. 7I, S7D). Thus, IL-1 signaling functions downstream of barrier dysregulation to regulate dermal γδT accumulation and the associated AD-like pathology.

**Figure 7.**
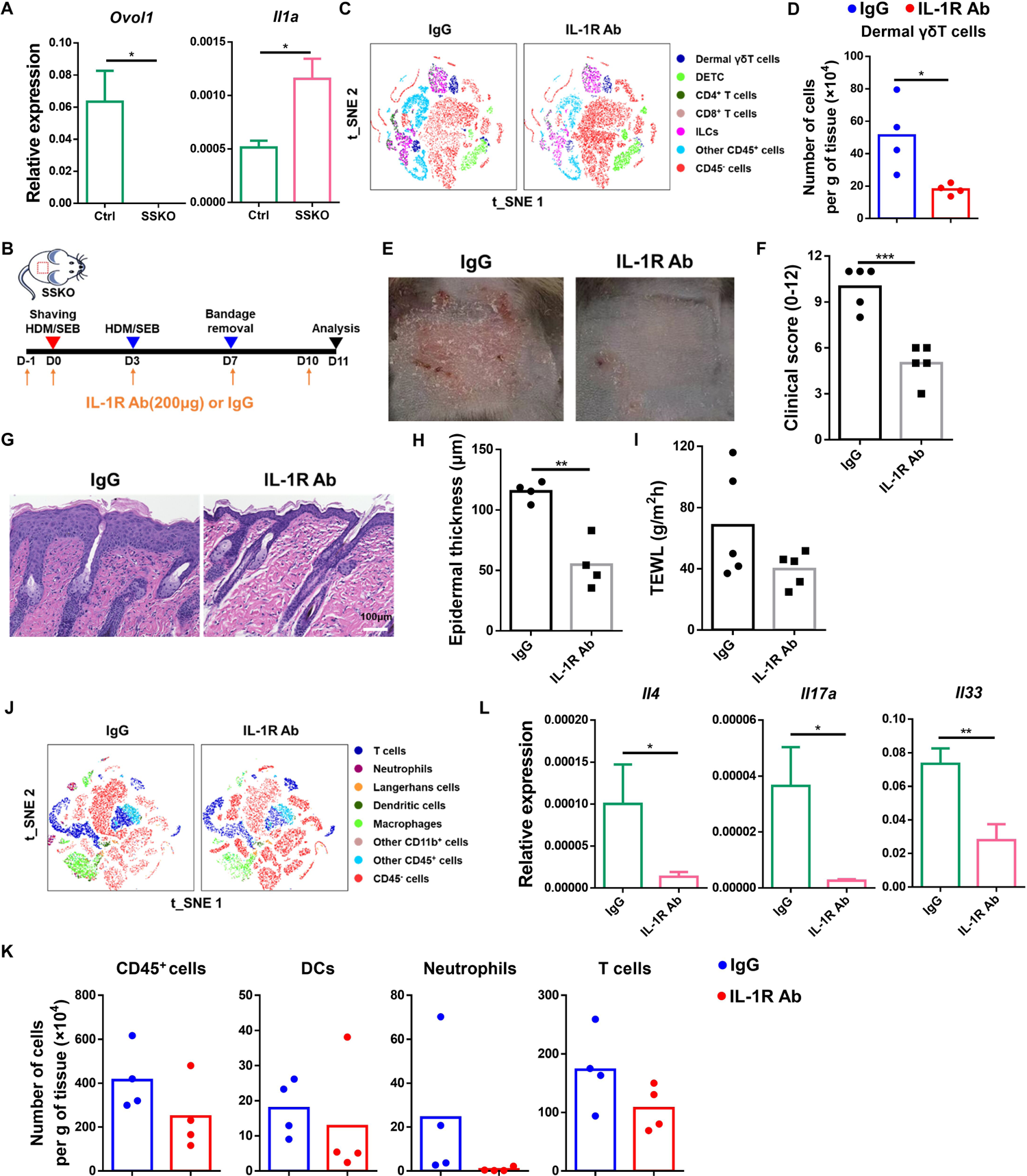
Blocking IL-1 signaling lessens dermal γδT abundance and skin inflammation in HDM/SEB-treated SSKO mice. **(A)** RT-qPCR results of *Ovol1* and *Il1a* expression in epidermal cells of the HDM/SEB-treated mice at day 7. n = 3 mice per group. **(B)** Experimental design for IL-1R Ab treatment in (C-K). Samples were harvested at day 11 after treatment for downstream analysis. IgG was used as a control. **(C-D)** Flow cytometry analysis of dermal γδT cells in the back skin of IgG- or IL-1R Ab-treated SSKO mice. See Fig. 5 legends for additional details. n = 4 mice per group. **(E)** Representative photographs of IgG- or IL-1R Ab-treated SSKO mice. **(F)** Clinical scores of IgG- or IL-1R Ab-treated SSKO mice. n = 5 mice per group. **(G)** Representative skin histology (H/E staining). Scale bar = 100 μm. **(H)** Quantification of epidermal, dermal and total skin thickness in IgG- or IL-1R Ab-treated SSKO mice. n = 4 mice per group. **(I)** TEWL values of IgG- or IL-1R Ab-treated SSKO mice. n = 5 mice per group. **(J-K)** Flow cytometry analysis of the indicated cell types in the back skin of IgG- or IL-1R Ab-treated SSKO mice. See Fig. 5 legends for additional details. n = 4 mice per group. **(L)** RT-qPCR results of *Il4*, *Il17a* and *Il33* expression in whole skin. Data are mean + SEM. n = 5 mice per group. * *p* < 0.05, ** *p* < 0.01, *** *p* < 0.001. *p* values were calculated using 2-tailed unpaired Student *t* test or Mann-Whitney U test.

We also comprehensively analyzed the impact of IL-1 signaling inhibition on other immune cell types. Reductions in IL-1 signaling-inhibited, HDM/SEB-treated SSKO mice relative to control (IgG) counterparts in most populations examined, including total immune cells, macrophages, LCs, DCs, ILCs, and other T cells, did not reach statistical significance (Fig. 7C, 7J-K, S7A-B, S7E). However, neutrophils, though of low abundance in SSKO skin, became barely detectable with IL-1R antibody treatment (Fig. 7J-K, S7A-B). The differential extent of changes in neutrophil abundance (3.9-fold and 31-fold decreases with γδTCR and IL-1R antibodies, respectively) implicates the existence of both γδT-dependent and γδT-independent actions of IL-1 signaling in promoting neutrophil accumulation. At a molecular level, the expression of *Il4, Il17a*, and *Il33*, but not *Tslp*, *Ccl20* and *Ccl27a,* was dramatically decreased in the skin lesions of IL-1R antibody-treated SSKO mice compared to IgG-treated SSKO littermates (Fig. 7L, S7F). These data suggest an important role for IL-1 signaling in instigating both type 2 and type 3 immune responses in AD-like SSKO skin.

A central, albeit not the only, regulatory theme that emerged from our data described so far is that epidermal loss of *Ovol1* results in barrier dysregulation and consequently IL-1 signaling enhancement, which augments the accumulation of dermal γδT cells that in turn orchestrate additional innate (neutrophil) and adaptive (DCs, Th2) immune responses in AD-like skin inflammation. To further dissect the link between epidermal barrier dysregulation and innate immune response, we next examined the immune impact of Id1 inhibition in SSKO AD skin. Flow cytometry analysis showed that Id1 inhibition drastically reduced the abundance of neutrophils, while the abundance of dermal γδT and other immune cells was not altered (Fig. S7G-I). Moreover, the expression of *Il1a*, *Il4* and *Il17a* was not altered by Id1 inhibition (Fig. S7J). The differential effects of Id1 inhibition and IL-1 signaling blockage in SSKO AD skin demonstrate the complexity of mechanisms that connect epidermal/barrier perturbations to innate immune (e.g., dermal γδT, neutrophil) responses, as well as implicate a crucial and rather specific role of the Ovol1-Id1 regulatory axis in controlling neutrophil accumulation in inflamed skin.

### OVOL1 downstream targets are upregulated in human AD skin

The functional significance of Ovol1 in suppressing AD-like mouse skin inflammation prompted us to investigate whether the expression of its target genes is altered in human AD skin. Cross-analysis of our Ovol1 ChIP-seq data with the published RNA-seq data of human AD epidermis (*42*) led to the identification of human homologs of 125 Ovol1 target genes that showed significantly increased expression in AD epidermis (Fig. 8A-B, S8, Table S15). These included *AQP3* and *ID1*, but not *KRT14,* again pointing to both molecular conservation and divergence between human and mouse skin. Immunostaining experiments also revealed the dramatically increased expression of AQP3 and ID1 proteins in AD epidermal keratinocytes (Fig. 8C-D). Thus, *Ovol1* deficiency-induced upregulation of *Aqp3* and *Id1* in mouse AD-like skin may be mirrored in the pathogenesis of human AD.

**Figure 8.**
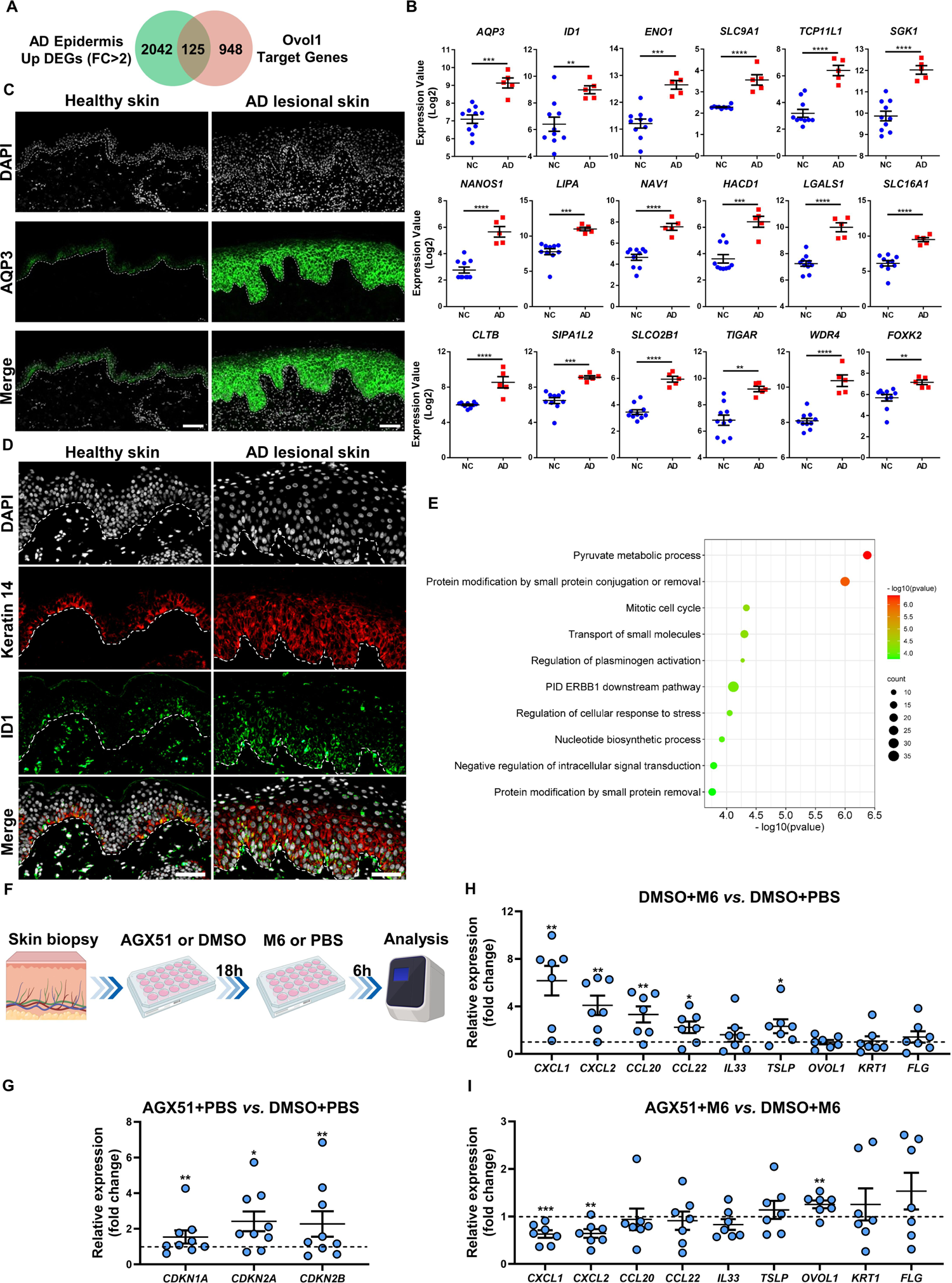
OVOL1 target gene expression in human AD skin. **(A)** Venn diagram of ChIP-seq-identified Ovol1 target genes and genes upregulated in human AD epidermis. The AD data were derived from published RNA-seq results (GSE120721). Gene numbers are indicated in the Venn diagram. **(B)** Expression of a select number of overlapping genes from (A) in the epidermis of normal (n = 10) and AD lesional (n = 5) skin. **(C)** Immunostaining of AQP3 protein in healthy and AD lesional skin. n = 4 samples per group. Dashed lines indicate the basement membrane. DAPI stains the nuclei. Scale bar = 100 μm. **(D)** Immunostaining of keratin 14 and ID1 proteins in healthy and AD lesional skin. n = 4 samples per group. Dashed lines indicate the basement membrane. DAPI stains the nuclei. Scale bar = 50 μm. **(E)** Ingenuity pathways analysis of the 125 overlapping genes in (A). **(F)** Experimental design for *ex vivo* human skin biopsy stimulation in (G-I). **(G)** RT-qPCR results of the indicated genes in AGX51- or DMSO-treated human skin biopsy with PBS treatment. n = 9 samples per group. **(H)** RT-qPCR results of the indicated genes in M6- or PBS-treated human skin biopsy with DMSO treatment. n = 7 samples per group. **(I)** RT-qPCR results of the indicated genes in AGX51- or DMSO-treated human skin biopsy with M6 treatment. n = 7 samples per group. ** *p* < 0.01, *** *p* < 0.001, **** *p* < 0.0001. *p* values were calculated using 2-tailed unpaired Student *t* test.

To gain further insights into the OVOL1-regulated events in human AD skin, we performed Metascape analysis on the 125 genes from above. This revealed “pyruvate metabolic process”, “protein modification by small protein conjugation or removal”, “mitotic cell cycle”, and “transport of small molecules” as top terms (Fig. 8E, Table S16). Pyruvate metabolism is known to promote keratinocyte proliferation (*66*). Genes included in this pathway and upregulated in AD skin are *ENO1*, *SLC9A1, FOXK2*, *LIPA*, *SLC16A1*, *TIGAR*, *ATF3*, *PPP4R3B*, *PEX13*, *HACD1*, *PANK2*, *GLDC*, *NDUFAF1*, *LANCL2*, *PRPS1*, *LPCAT3*, *DTYMK*, *PATL1*, *CD44* and *PFKP* (Fig. 8B).

*ENO1*-encoding enolase-1 is a glycolytic enzyme that has been reported to disrupt the tight junction barrier in AD keratinocytes (*67*). *SLC9A1* encodes sodium–hydrogen antiporter 1, a membrane-bound enzyme involved in volume- and pH-regulation, which is known to regulate stratum corneum permeability barrier homeostasis (*68*). *FOXK2* is a transcription factor that reprograms cellular metabolism to induce aerobic glycolysis (*69*). *LIPA* encodes a lysosomal acid lipase that hydrolyzes cholesteryl esters and triglycerides (*70*). *SLC16A1* encodes monocarboxylate transporter 1, which transports monocarboxylates such as pyruvate and lactate into the cells, and lactate is known to regulate hair follicle stem cell activation (*71*). *TIGAR*, TP53-inducible glycolysis and apoptosis regulator, encodes an enzyme that regulates glycolysis and ROS production (*72*). Thus, our data implicate a previously unrecognized role of OVOL1 in regulating epidermal cellular metabolism and barrier with potential relevance to human AD pathogenesis.

Finally, as proof-of-principle validation of the functionality of OVOL1 targets, we tested the effect of ID1 inhibitor AGX51 on explant cultures derived from skin biopsies of healthy individuals (Fig. 8F). We found AGX51 treatment to significantly increase the expression of several canonical ID1 target genes, including cell cycle regulators *CDKN1A*, *CDKN2A,* and *CDKN2B* (*73*) (Fig. 8G). We also treated the explants with a cytokine cocktail (including IL-17A, IL-22, oncostatin M, IL-1α, and TNF-α) known to stimulate cultured keratinocytes to produce multiple cytokines and chemokines related to human skin inflammation (*74*). This approach, combined with the inclusion of IL-4 (the resulting cocktail is called M6), was employed to establish an *ex vivo* human AD-like skin model (Fig. 8F). We found that M6 treatment alone resulted in elevated expression of *CXCL1*, *CXCL2*, *CCL20*, *CCL22* and *TSLP* (Fig. 8H), whereas the addition of AGX51 dampened this effect, evident through the significantly reduced expression of neutrophil chemoattractant *CXCL1* and *CXCL2* (Fig. 8I). Intriguingly, AGX51 treatment also slightly but significantly increased *OVOL1* expression in M6-treated skin explants, which may reflect altered differentiation or possible feedback regulation. Taken together, our data show that elevated activity of OVOL1 target ID1 has the capacity to functionally contribute to human skin inflammation.

## DISCUSSION

Proper regulation of epidermal keratinocyte proliferation and differentiation is integral to the maintenance of a robust permeability barrier especially under external challenges. Keratinocytes are also known to secrete cytokines and chemokines that stimulate immune cell activities to drive the pathogenesis of inflammatory skin diseases such as AD (*4*). Our study has now uncovered an OVOL1/Ovol1-directed gene expression program that operates within keratinocytes and directly downstream of AhR to suppress cell cycle and proliferation, regulate cellular metabolism, promote terminal differentiation and barrier integrity, as well as modulate host immune defense against AD-inducing allergen and pathogen (Fig. 9). As such, our discoveries unveil an intricate transcriptional regulatory mechanism that coordinates the homeostasis of both the epidermis and the immune system at the barrier interface against assaults leading to AD.

**Figure 9.**
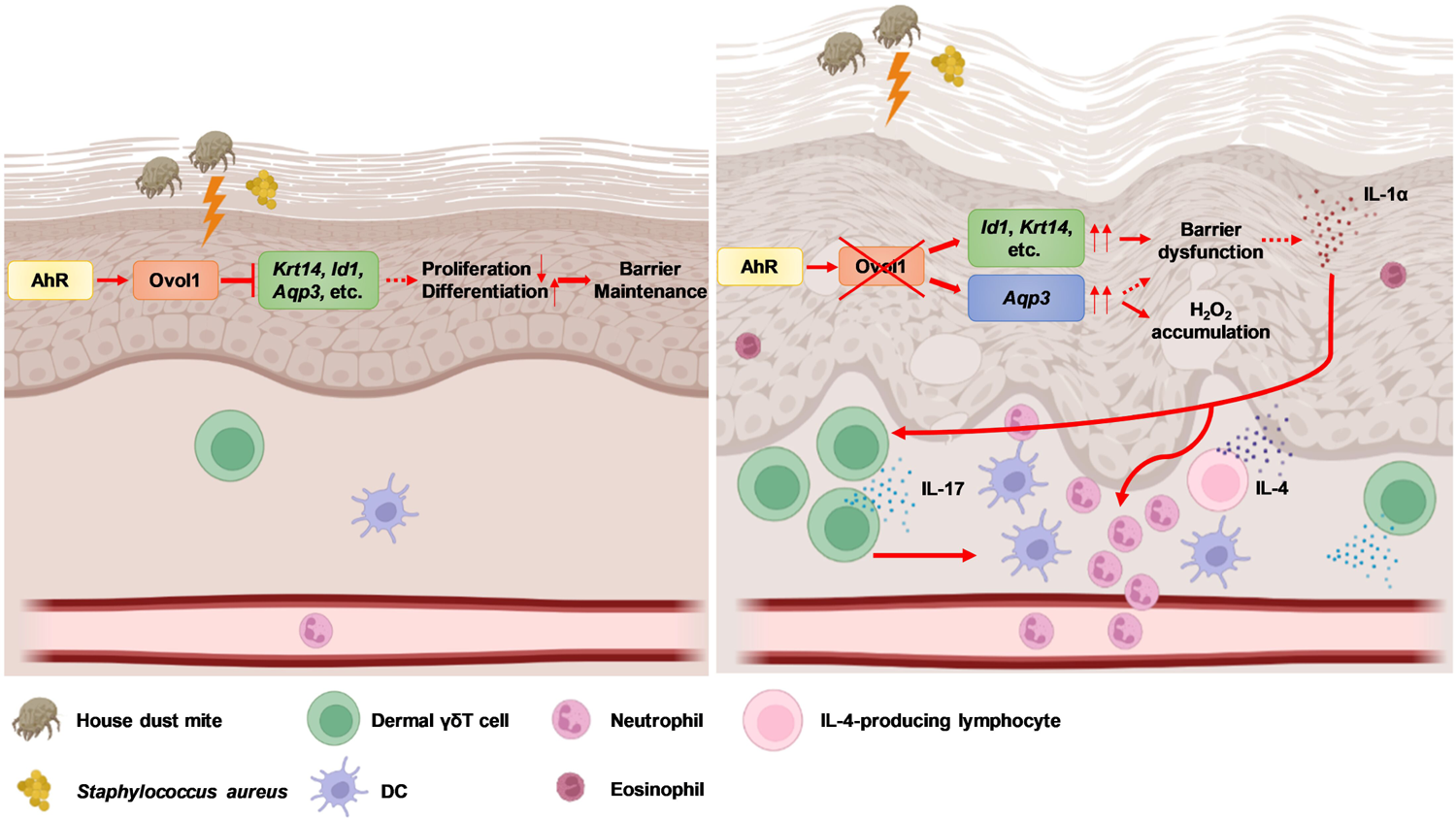
Working model on Ovol1 regulation of epidermal and immune homeostasis during AD-like skin inflammation. Solid lines indicate regulation for which we show evidence in this study. Dashed lines indicate regulation reported in the literature. Thin upward arrows indicate aberrant upregulation when *Ovol1* is ablated. This figure was created using BioRender.com.

AhR agonist Tapinarof shows potential and/or efficacy in treating both AD and psoriasis, consistent with a role in targeting the early and common events of skin inflammation. However, the underlying mechanisms of action remain largely unclear, and AhR activation is known to elicit both anti-inflammatory and pro-inflammatory consequences (*75, 76*). As such, dissecting the downstream mediators, especially the anti-inflammatory arm, of AhR function in skin is important from both basic science and clinical perspectives. Our finding of *OVOL1*/*Ovol1* as a direct target and key mediator of the barrier-protecting/anti-inflammatory function of AhR now lays the ground work for targeting the OVOL1-associated cellular and molecular components in order to find therapeutics that effectively prevent and treat inflammatory skin diseases but with minimal adverse effects.

Our work also implicates new players in skin inflammation and AD pathogenesis. In particular, our human and mouse studies collectively suggest a role for ID1/Id1, a direct target of OVOL1/Ovol1 repression in keratinocytes, in promoting AD-associated epidermal hyperplasia, barrier disruption, inflammation, and clinical symptoms. While ID1 is known to be required for keratinocyte proliferation (*51*), its role in skin inflammation has not been previously reported. The relief of neutrophil accumulation, but not abundance of other immune cells, by inhibition of Id1 in *Ovol1*-deficient AD-like skin raises an interesting possibility that Id1 may regulate neutrophil infiltration in a more specific manner than simply augmenting barrier dysregulation-associated immune consequences. Our *ex vivo* experiments with human skin biopsies showing the specific effect of ID1 inhibition on the expression of neutrophil chemoattractants, *CXCL1* and *CXCL2*, support this notion. This said, we cannot formally exclude the possibility that systemic administration of AGX51 to SSKO mice reduces neutrophil infiltration in part through affecting neutrophil development and maturation (*77*).

Using *Ovol1* SSKO mice as a model, we have obtained fundamental insights into how dermal γδT cells are regulated in, and how they shape, AD-like skin inflammation. Dermal γδT cells are a subset of innate T cells that can respond rapidly to intrinsic or extrinsic stimuli (*78*). Skin barrier damage after tape striping induces IL-22 expression in dermal γδT cells and this process depends on keratinocyte-derived IL-23 (*79, 80*). Neither *Il23* nor *Il22* expression is upregulated in HDM/SEB-treated SSKO skin. Instead, our data show barrier dysregulation-induced IL-1 signaling to be critical for amplifying the dermal γδT cell response during AD-like skin inflammation in *Ovol1*-deficient mice. Furthermore, our finding of *Aqp3* as a direct target of Ovol1 with increased expression in *Ovol1*-deficient AD-like skin provides an additional, likely more direct connection between *Ovol1* deletion in the epidermis and γδT cell expansion in the dermis, because *Aqp3* is recognized for facilitating the accumulation of γδT cells in psoriatic mouse skin (*48*). There are two major subsets of dermal γδT cells, Vγ2^+^ and Vγ4^+^ in murine skin (*78*). Both subsets are pro-inflammatory in psoriasis-like skin inflammation (*78*), whereas their roles in spontaneous AD-like skin inflammation is controversial (*81, 82*). We indeed detected two subsets of dermal γδT cells in our flow analysis, and both subsets showed elevated abundance in *Ovol1* SSKO AD-like skin (Fig. 5B). Furthermore, our antibody blocking data support a pro-inflammatory function of dermal γδT cells in HDM/SEB-induced AD-like inflammation.

We find that γδTCR signaling is required not only for maximal neutrophil infiltration, as previously shown in non-AD contexts such as contact dermatitis and cutaneous bacterial infection (*30, 31, 78*), but also for DC expansion in the AD-like skin. This latter effect is unlikely through an IL-17A/Ccl20 axis (*83, 84*), because γδTCR antibody treatment did not decrease *Ccl20* expression in skin lesions of HDM/SEB-treated SSKO mice. More importantly, both functional blocking of γδT cells and inhibition of IL-1 signaling result in reduced expression of not only *Il17* but also type 2 cytokine *Il4*, providing evidence for a role of the IL-1/γδT axis in orchestrating type 2 immunity. As DCs are known recruiters of both type 2 (Th2) and type 3 (Th17) T cells to inflamed skin (*4*), γδT-facilitated DC accumulation could potentially be a mechanism by which Th2 cells are recruited to *Ovol1* SSKO AD-like skin. Interestingly, DCs are known to activate dermal γδT cells (*83*). Our data, in conjunction with this observation, highlights the reciprocal interaction between innate and adaptive immune components, collectively driving skin inflammation.

Although γδT cells are rarely detected in human skin (*85*), the number of peripheral γδT cells positively correlates with AD disease severity (*86*). Additionally, akin to Th17 cells, dermal γδT cells produce IL-17A and exhibit similar responses to cytokines and chemokines (*65, 83*). Therefore, the insights gained from our study on the regulation of dermal γδT cells by the epidermis in mouse skin may have translational implications for understanding epidermal-Th17 interactions in human skin during the early stages of disease progression and/or in specific subtypes of AD (*15*). Along the same line, while not all molecular intricacies of the AhR-Ovol1-Id1 axis are identical between human and mouse epidermis, the fundamental regulations in, and functions of, this axis appear to be conserved. By deciphering the activation status of AhR and ensuring proper keratinocyte differentiation to bolster a resilient epidermal barrier, Ovol1 in keratinocytes effectively orchestrates the maintenance of both epidermal and immune homeostasis, thereby preventing disease exacerbation.

## MATERIALS AND METHODS

### Cell culture and transfection

NHEKs were cultured in Keratinocyte Serum-Free Medium (Gibco, Cat# 17005042) per manufacturer’s protocol. NHEKs within passage 3-6 were used for experiments. Scrambled, *AHR*- or *OVOL1*-specific siRNA (generated by Huzhou Hippo Biotechnology Co., Ltd) was transfected into NHEKs using Lipofectamine RNAiMAX Transfection Reagent (Thermo Fisher Scientific, Cat# 13778075) per manufacturer’s protocol.

### RNA-seq and RT-qPCR

For bulk RNA-seq, NHEKs were transfected with scrambled or *OVOL1*–specific siRNA for 24 hours, and then treated with 100 nM FICZ (MedChemExpress, Cat# HY-12451) or 10% v/v of TSB (100 μl TSB to 900 μl medium, Sigma-Aldrich, Cat# 22090) for another 24 hours. Total RNA was extracted using Trizol reagent (Invitrogen, Waltham, MA). Library preparation, sequencing and analysis of differentially expressed RNAs were performed as described previously (*87*). DEGs were identified using *p* value < 0.05 and >1.5-fold change as cut-offs. Enriched pathways of DEGs were analyzed using Metascape (*88*).

Mouse back skin was homogenized and RNA was extracted using Quick-RNA Miniprep Kit (Zymo Research, Cat# R1054) per the manufacturer’s instructions. For RT-qPCR, 1 μg of RNA was used to generate cDNA (Applied Biosystems, Cat# 4368814) per manufacturer’s protocol. RT-qPCR was performed using a Bio-Rad CFX96 Real-Time System and SsoAdvanced Universal SYBR® Green Supermix (Bio-Rad, Cat# 172-5271). Information for gene-specific primers used is provided in Table S17. Public RNA-seq data of primary keratinocytes from *Ahr*^+/+^ and *Ahr*^-/-^ mice (GSE62490) (*37*) and epidermis from normal and AD human skin (GSE120721) (*89*) were analyzed using the online tool GEO2R. Human single-cell RNA-seq data were downloaded from https://zenodo.org/record/4310074#.Yd4yDdBBxPd and analyzed using Python as described previously (*44*).

### Mice

*K14-*Cre;*Ovol1*^+/-^ (C57BL/6 background), *Ovol1*^-/-^ (CD1 background), and *Ovol1*^flox/flox^ (C57BL/6 background) mice were generated as described previously (*30, 31*). Sex- and weight-matched control and mutant littermates were used for all experiments. Both male and female mice at 7-10 weeks of age were used. All animal studies have been approved and abide by regulatory guidelines of the Institutional Animal Care and Use Committee of the University of California, Irvine.

### AD models

Mice were epicutaneously sensitized with HDM and SEB as described previously (*43*). In brief, a solution of 10 μg HDM extract (Stallergen Greer) and 500 ng SEB (List Biological Laboratories) in PBS was applied on a 1 cm^2^ gauze pad placed on the shaved skin and occluded with a Tegaderm Transparent Dressing (3 M HealthCare). Three days later, the gauze pads were replaced. Four days later, dressings were removed and mice were kept without treatment for 4 days and then sacrificed for analysis. For experiments with three rounds of treatment, mice received this “3+4-d” pattern of treatment twice more and were sacrificed two days after the last cycle of treatment. TEWL was measured on shaved mouse back skin using the Delfin VapoMeter (SWL4400). TEWL values are output as g/m^2^h.

Two regimens were employed for the MC903-induced AD model. For *Ovol1*^-/-^ mice, we topically applied 4 nmols of MC903 (Sigma, St.Louis, Missouri) or ethanol control onto the mouse ears once daily for 14 consecutive days. Mice were sacrificed at day 15. For SSKO mice, we topically applied 1.125 nmols of MC903 or ethanol onto the mouse ears once daily for 5 consecutive days; mice were then kept without treatment for 2 days, followed by topical application of 1.125 nmols of MC903 once daily for 3 more days. Mice were sacrificed at day 11. Ear thickness was measured using Caliper.

For *ex vivo* human AD model, skin tissues (approximately 0.25 or 0.5 cm^2^) from 9 healthy individuals were obtained by excisional biopsy during plastic surgery and divided into 2 or 4 equal parts. The tissues were cultured in DMEM medium (Gibco, Cat# 11965092) containing 10% FBS and treated with AGX51 (100 μM) or DMSO for 18 hours, followed by treatment with M6 cytokine cocktail or PBS for another 6 hours. The M6 cocktail contains IL-4 (10 ng/ml, Peprotech, Cat# 200-04), IL-17A (10 ng/ml, Peprotech, Cat# 200-17), IL-22 (10 ng/ml, Peprotech, Cat# 200-22), oncostatin M (10 ng/ml, Peprotech, Cat# 300-10), IL-1α (10 ng/ml, Peprotech, Cat# 200-01A), and TNF-α (10 ng/ml, Peprotech, Cat# 300-01A). After M6 treatment, skin tissues were collected and stored in liquid nitrogen until further analysis.

### *In vivo* FICZ, antibody, and AGX51 treatment

For FICZ experiments, mice were i.p injected with 100 μg/kg FICZ (MedChemExpress, Cat# HY-12451) or DMSO in 100 μl corn oil once daily from day 0 to day 10. For γδTCR blocking, mice were i.p injected with 500 μg γδTCR antibody (Biolegend, Cat# 107517, Clone UC7-13D5) or IgG (Biolegend, Cat# 400959, Clone HTK888) at day 0, followed by 200 μg at days 3, 7 and 10 of HDM/SEB treatment. For IL-1R blocking, mice were i.p injected with 200 μg IL-1R antibody (BioXcell, Cat# BE0256, Clone JAMA-147) or IgG (BioXcell, Cat# BE0091) at days −1, 0, 3, 7 and 10 of HDM/SEB treatment. For AGX51 experiments, mice were i.p injected with 1 mg AGX51 (MedChemExpress, Cat# HY-129241) or DMSO in 100 μl corn oil once daily from day 0 to day 10.

### Histology, indirect immunofluorescence, and RNAScope

Sections from paraformaldehyde-fixed, paraffin-embedded back skin were stained with hematoxylin and eosin (H/E) as previously described (*30*). Epidermal and dermal thickness was measured using ImageJ. For indirect immunofluorescence, mouse back skin tissues were freshly frozen in optimum cutting temperature (OCT) compound (Tissue-Tek) and stained using the appropriate antibodies as described previously (*30*). The primary antibodies used were: keratin 14 and filaggrin (rabbit or chicken, gifts of Julie Segre, National Institutes of Health, Bethesda), Ki67 (rabbit, Cell Signaling Technology, Cat# 9129, Clone D3B5), Ly6G (rat, eBioscience, Cat# 16-9668-82, Clone 1A8). RNAScope was performed as described (*90*) using *Krt14* probe (ACD, Cat# 422521-C3).

### Patient sample analysis

Skin samples were obtained from 4 AD patients and 4 healthy individuals by excisional biopsy. Sample acquisitions, including skin biopsies, were approved by the Ethics Committee of Shanghai Tenth People’s Hospital and performed in accordance with the declaration of Helsinki Principles. Informed consent was obtained for all procedures.

Paraffin blocks of skin samples were used to obtain 4-μm sections, which were used for immunofluorescence and fluorescence in situ hybridization as previously described (*91*). Antibodies used for immunofluorescence were anti-cytokeratin 10 (rabbit, Servicebio, Cat# GB112105-100), anti-cytokeratin 14 (rabbit, Abcam, Cat# ab119695), anti-ID1 (mouse, Santa Cruz, Cat# sc-133104), and anti-aquaporin 3 (rabbit, Servicebio, Cat# GB11395-100). Immunostaining of AQP3 or co-staining of keratin 14 and ID1 was performed using a Three-color Fluorescence kit (Recordbio Biological Technology, Cat# RC0086-23RM) based on the tyramide signal amplification technology according to the manufacture’s instruction. For maximum detection of *OVOL1* RNA, a 1:1:1:1 mixture of four different probes (Table S17) was used.

### Western blotting

HaCaT cells were treated with 100 μM AGX51 or DMSO for 24 hours. Protein extraction and Western blotting were performed as described previously (*92*). ID1 antibody (Genetex, Cat# GTX133738, 1:1000) was used.

### Single cell suspension and flow cytometry

Single cell suspension of skin was prepared as described previously (*30*). In brief, to obtain epidermal cell suspension, subcutaneous fat tissue was removed from the skin and skin was treated with 0.25% trypsin with the dermis side facing down for 45 min at 37℃. Epidermis was then separated from dermis, minced using a fine scissor and filtered through a 40 μm filter. To obtain whole skin cell suspension, skin was minced using a fine scissor and then digested with 10 ml of RPMI 1640 Medium (Gibco, Cat# 11875093) containing 0.25% collagenase (Sigma, Cat# C9091), 0.01 M HEPES (Thermo Fisher Scientific, Cat# BP310), 0.001 M sodium pyruvate (Thermo Fisher Scientific, Cat# BP356), and 0.1 mg/ml DNase (Sigma, Cat# DN25) for 1-1.5 hours at 37 ℃ with rotation, and then filtered through a 40 μm filter. Cells were spun down and resuspended in FACS buffer (2% fetal bovine serum in PBS). For cell surface staining, cells were incubated with TruStain FcX™ Antibody (Biolegend, Cat# 101319) to block Fc receptors for 10 min at room temperature and incubated with antibodies for 30 min at 4 ℃. For intracellular staining, cells were incubated with Zombie NIR™ Fixable Viability Kit (Biolegend, Cat# 423105) for 30 min at 4 ℃ before cell surface staining. After cell surface staining, cells were fixed with IC fixation buffer (eBioscience, Cat# 00-8222-49) for 30 min at 4 ℃, then stained with antibodies in permeabilization buffer (eBioscience, Cat# 00-8333-56) for 30 min at 4 ℃. Cells with surface staining only were incubated with SYTOX™ Blue Dead Cell Stain (Invitrogen, Cat# S34857) for 5 min before analysis. Cells were analyzed using BD FACSAria Fusion Sorter. Data were analyzed using FlowJo v10.7.2. Antibodies used were: Brilliant Violet 510 anti-mouse CD45 (Biolegend, Cat# 103137, Clone 30-F11), allophycocyanin (APC) anti-mouse CD45 (Biolegend, Cat# 103112, Clone 30-F11), Alexa Fluor 488 anti-mouse CD11b (Biolegend, Cat# 101219, Clone M1/70), APC/Cyanine7 anti-mouse Ly6g (Tonbo Biosciences, Cat# 25-1276, Clone 1A8), APC anti-mouse CD11c (Biolegend, Cat# 117310, Clone N418), phycoerythrin (PE) anti-mouse F4/80 (Biolegend, Cat# 101219, Clone BM8), PE/Cyanine7 anti-mouse EpCAM (Biolegend, Cat# 118216, Clone G8.8), PerCP/Cyanine5.5 anti-mouse CD3 (Biolegend, Cat# 100217, Clone 17A2), Alexa Fluor 700 anti-mouse lineage (Biolegend, Cat# 133313), PE anti-mouse γδTCR (Biolegend, Cat# 118108, Clone GL3), Brilliant Violet 605 anti-mouse CD4 (Biolegend, Cat# 100548, Clone RM4-5), Fluorescein isothiocyanate (FITC) anti-mouse CD8 (Biolegend, Cat# 100706, Clone 53-6.7), PerCP/Cyanine5.5 anti-mouse CD90.2 (Biolegend, Cat# 105338, Clone 30-H12), FITC anti-mouse MHC II (Invitrogen, Cat# 11-5321-82, Clone M5/114.15.2), and PE anti-mouse CD207 (Invitrogen, Cat# 12-2075-82, Clone eBioL31).

### ChIP analysis

Mouse primary keratinocytes were treated with CaCl_2_ (1.8 mM) for 24 hours and then cross-linked in 1% formaldehyde. NHEKs were treated with FICZ (100 nM) for 3 hours and then cross-linked in 1% formaldehyde. ChIP assay was performed using anti-OVOL1 antibody (rabbit, GeneTex, Cat# GTX55272), anti-AhR antibody (rabbit, Cell Signaling Technology, Cat# 83200S), and SimpleChIP® Enzymatic Chromatin IP Kit (Agarose Beads) (Cell Signaling Technology, Cat# 9002) according to manufacturer’s instructions. DNA was then purified and RT-qPCR performed. Information for gene-specific primers used is provided in Table S16.

Ovol1 ChIP-seq data were analyzed as described previously (*32*). Peaks were called with MACS2 broad peak calling (*93*). Motifs were analyzed using HOMER v4.11 (*94*). Public ChIP-seq BIGWIG data were downloaded from GEO datasets and visualized using IGV_2.10.2. The ChIP-seq data used were: GSE86900 (Mouse H3K4me1, H3K4me3 and H3K27ac), GSE72455 (Mouse AhR), GSM733674 (NHEK H3K27ac), GSM733698 (NHEK H3K4me1), GSM733720 (NHEK H3K4me3) and GSM733740 (NHEK Input).

### Intracellular H_2_O_2_ measurement

Intracellular level of H_2_O_2_ was measured as described previously (*48*). In brief, epidermal cells from mouse back skin were incubated with H2DCFDA (Invitrogen, Cat# D399) at a concentration of 10 μM at 37℃ for 30 min. Cells were then analyzed using BD FACSAria Fusion Sorter, and data analyzed using FlowJo v10.7.2.

### Measurement of serum levels of Cxcl2 and G-CSF

Custom Plex Assays (Eve Technologies) were used to detect the serum levels of Cxcl2 and G-CSF in MC903-treated mice according to manufacturer’s instructions.

### Statistical analysis

Two-tailed unpaired Student *t* test was used for comparing two groups if the data were normally distributed. Mann-Whitney U test was used for comparing two groups if the data were not normally distributed. Two-way analysis of variance (ANOVA) was used for comparing multiple groups. All analyses except single-cell RNA-seq data were performed using Prism 6 (GraphPad). *P* values of single-cell RNA-seq data were calculated using T test followed by Benjamini-Hochberg correction.

## ACKNOWLEDGEMENT

We thank the Institute for Immunology FACS Core Facility at the University of California, Irvine for expert service, Bogi Andersen for use of the Vapometer, Scott Atwood for NHEKs, Xilin Zhang and Bingjie Li for advice and support.

## Funding

Z.C. received support from the China Scholarship Council (Grant No. 201906260233). M.D., D.H., and R. V. received support from National Science Foundation (NSF)-Simons Predoctoral Fellowship (NSF grant DMS1763272 and Simons Foundation grant 594598). M.D. was supported by the NIH T32 Immunology Research Training Grant (AI 060573). R.V. was supported by NSF predoctoral fellowship DGE-1839285. Y.S. was supported by grants from National Natural Science Foundation of China (No. 81872522, 82073429) and Innovation Program of Shanghai Municipal Education Commission (No.2019-01-07-00-07-E00046). X.D. was supported by National Institutes of Health Grants R01-AR068074, R35-GM145307, U01-AR073159, P30-AR075047, NSF grant DMS1763272, Simons Foundation grant 594598, and UCI Office of Research.

### Author contributions

Conceptualization: X.D.; Methodology: Z.C., M.D., D.H., R.V., L.C., P.S.; Investigation: Z.C., M.D., D.H., R.V., L.C., P.S.; Visualization: Z.C., M.D., D.H., R.V.; Funding acquisition: X.D., Y.S.; Project administration: X.D.; Supervision: X.D., Y.S.; Writing – original draft: X.D., Z.C.; Writing – review & editing: X.D., Z.C., M.D., P.S., D.H., R.V., L.C., and Y.S.

## Data and materials availability

All data needed to evaluate the conclusions in the paper are present in the paper or the Supplementary Materials. Request for materials and correspondence should be addressed to X.D. and Y.S.

## SUPPLEMENTAL TABLES

**Table S1.** List of the differentially expressed genes in NHEKs treated with siNC+FICZ *versus* siNC+DMSO.

**Table S2.** Metascape result of the FICZ-induced genes.

**Table S3.** Metascape result of the FICZ-suppressed genes.

**Table S4.** List of the differentially expressed genes in NHEKs treated with si*OVOL1*+DMSO *versus* siNC+DMSO.

**Table S5.** Metascape result of the downregulated genes upon *OVOL1* depletion in DMSO-treated NHEKs.

**Table S6.** Metascape result of the up-regulated genes upon *OVOL1* depletion in DMSO-treated NHEKs.

**Table S7.** List of the differentially expressed genes in NHEKs treated with si*OVOL1*+FICZ *versus* siNC+FICZ.

**Table S8.** Metascape result of the downregulated genes upon *OVOL1* depletion in FICZ-treated NHEKs.

**Table S9.** Metascape result of the up-regulated genes upon *OVOL1* depletion in FICZ-treated NHEKs.

**Table S10.** List of FICZ-induced or FICZ-suppressed and OVOL1-dependent genes.

**Table S11.** Metascape result of FICZ-induced and OVOL1-dependent genes.

**Table S12.** Metascape result of FICZ-suppressed and OVOL1-dependent genes.

**Table S13.** Annotated peaks of Ovol1 ChIP-seq analysis.

**Table S14.** Metascape result of Ovol1 target genes.

**Table S15.** List of the 125 human homologs of Ovol1 target genes that are upregulated in AD epidermis.

**Table S16.** Metascape result of the 125 human homologs of Ovol1 target genes that are upregulated in AD epidermis.

**Table S17.** Primer information.

## SUPPLEMENTAL FIGURE LEGENDS

**Figure S1.**
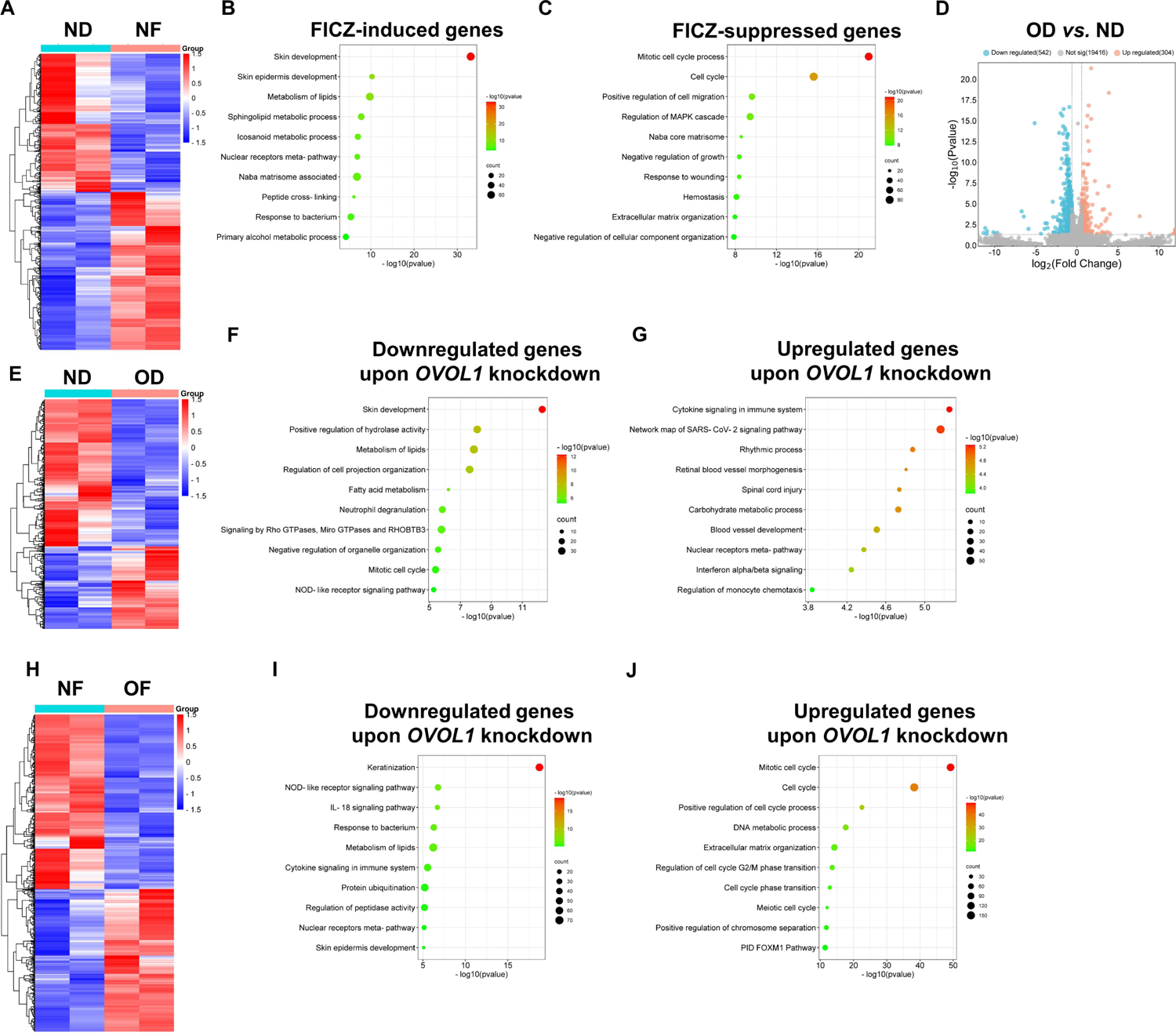
Metascape data of FICZ- or OVOL1-regulated genes in NHEKs. Related to Figure 1. **(A)** Heatmap of the DEGs between NF and ND. **(B-C)** Ingenuity pathways analysis of the FICZ-induced (B) or FICZ-suppressed (C) genes. **(D)** Volcano plots showing DEGs between OD and ND. n = 2 per group. **(E)** Heatmap of the DEGs between OD and ND. **(F-G)** Ingenuity pathways analysis of the downregulated (F) or upregulated (G) genes upon *OVOL1* depletion in DMSO-treated NHEKs. **(H)** Heatmap of the DEGs between OF and NF. **(I-J)** Ingenuity pathways analysis of the downregulated (I) or upregulated (J) genes upon *OVOL1* depletion in FICZ-treated NHEKs. ND: siNC + DMSO, NF: siNC + FICZ, OD: si*OVOL1* + DMSO, OF: si*OVOL1* + FICZ.

**Figure S2.**
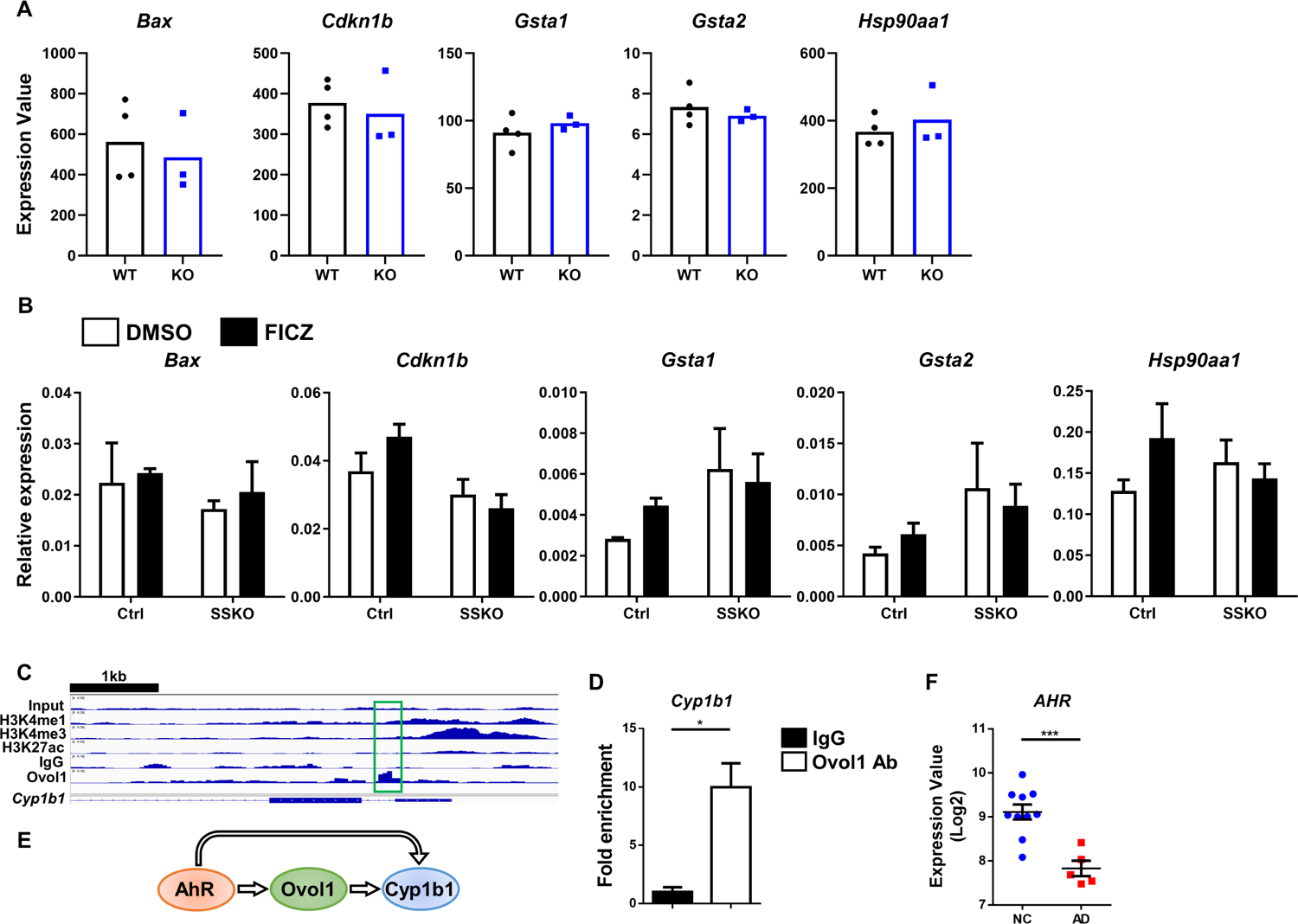
Impact of *Ahr* or *Ovol1* deletion on the expression of previously reported AhR targets. Related to **Figure 2. (A)** RNA-seq results for the indicated genes in primary keratinocytes from *Ahr*^+/+^ (WT, n = 4) or *Ahr*^-/-^ (KO, n = 3) mice. **(B)** RT-qPCR results of the indicated genes in the whole skin of control (Ctrl) and SSKO mice at day 11 after FICZ or DMSO treatment. Data are mean + SEM. n = 4 mice per group. **(C)** Genome browser track for the indicated ChIP-seq signals across the *Cyp1b1* locus. Green box highlights the Ovol1-bound region. **(D)** ChIP-qPCR results of *Cyp1b1* in mouse keratinocytes treated with Ca^2+^ (1.8 mM) for 24 hours. IgG control values were normalized to 1. Data are mean + SEM. Results are summarized from 3 independent experiments. **(E)** Working model of the regulation of *Cyp1b1* by AhR and Ovol1. **(F)** RNA-seq results (GSE120721) of *AHR* expression in the epidermis of healthy (NC, n = 10) and AD lesional (n = 5) human skin. * *p* < 0.05, *** *p* < 0.001. *p* values were calculated using 2-tailed unpaired Student *t* test.

**Figure S3.**
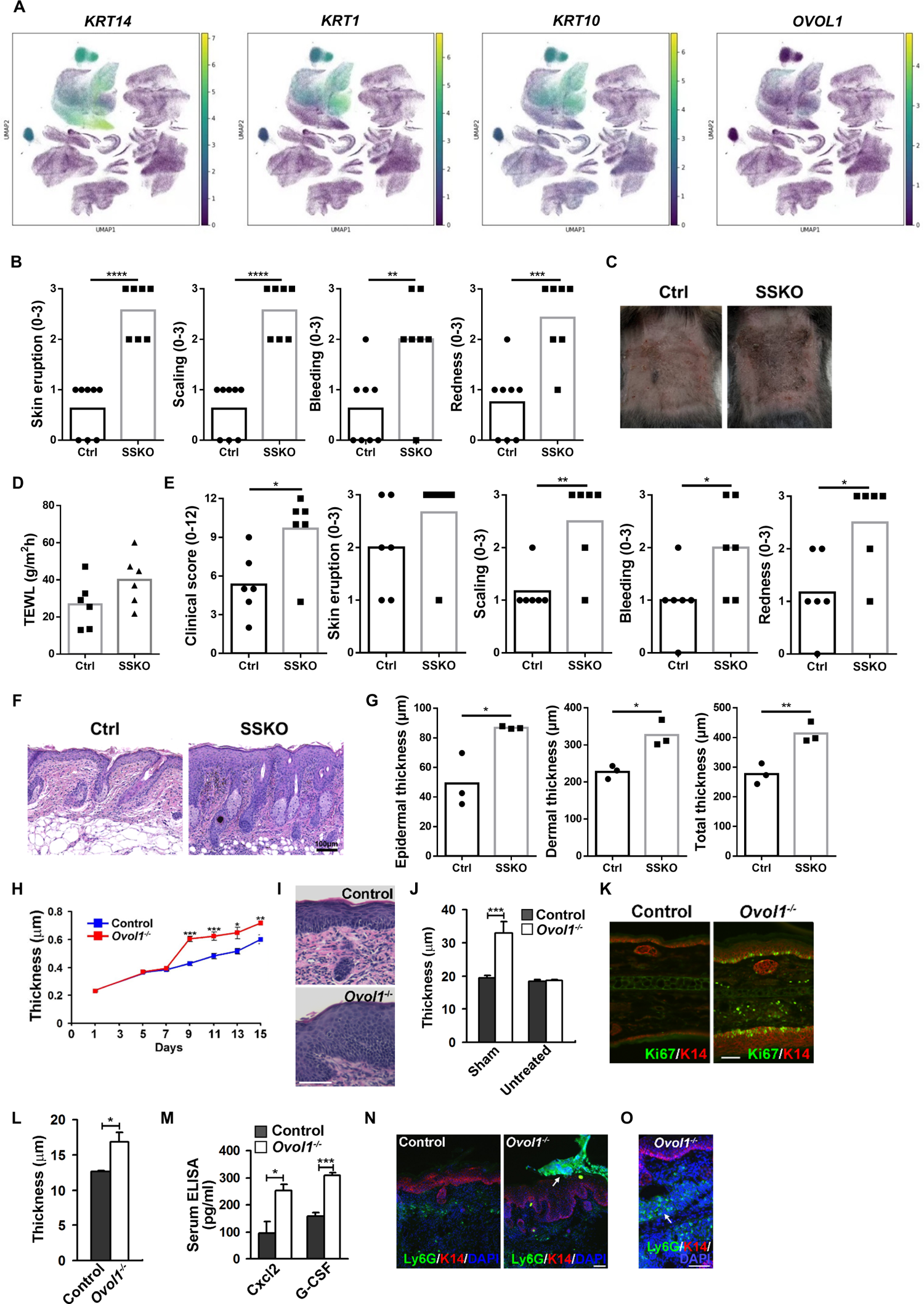
*Ovol1* deficiency in keratinocytes aggravates AD-like skin inflammation in mice. Related to **Figure 3. (A)** Feature plots showing expression of the indicated genes in all cells of healthy and AD lesional and non-lesional skin. **(B)** Clinical score of skin eruption, scaling, bleeding and redness in HDM/SEB-treated Ctrl (n = 8) and SSKO (n = 7) mice at day 11. **(C)** Representative photographs of HDM/SEB-treated Ctrl and SSKO mice at day 31. **(D)** TEWL values of HDM/SEB-treated Ctrl and SSKO mice at day 31. n = 6 mice per group. **(E)** Clinical scores of HDM/SEB-treated Ctrl and SSKO mice at day 31. n = 6 mice per group. **(F)** Representative skin histology (H/E staining) HDM/SEB-treated Ctrl and SSKO mice at day 31. Scale bar = 100 μm. **(G)** Quantification of epidermal, dermal and total skin thickness. n = 3 mice per group. **(H)** Mean ± SEM ear thickness of MC903-treated control (Ctrl) and *Ovol1*^-/-^ mice. n = 6 mice per group. **(I)** Representative skin histology of MC903-treated Ctrl and *Ovol1*^-/-^ mice at day 15. Scale bar = 100 μm. **(J)** Thickness of sham (ethanol-treated) ear skin at 9 days after MC903 application to the other ear of the same mouse (n = 6 per group) or in untreated (n = 3 per group) mice. **(K)** Ki67 and K14 immunostaining of sham-treated, left ears of mice that were treated with MC903 on their right ears. Scale bar = 50 µm. **(L)** Thickness of back skin epidermis in mice that were treated with MC903 on their right ears. n = 3 mice per group. **(M)** Serum ELISA analysis of mice that were treated with MC903 on their right ears. n = 3 mice per group. **(N-O)** Ly6G and K14 immunostaining of the MC903-treated right ears (N) and ethanol-treated (O) left ears. White arrows point to neutrophils that are present on the skin surface (N) or in dermis (O). DAPI stains the nuclei. Scale bar = 50 µm. * *p* < 0.05, ** *p* < 0.01, *** *p* < 0.001, **** *p* < 0.0001. *p* values were calculated using two-way ANOVA (H) or 2-tailed unpaired Student *t* test (B, D, E, G, J, L and M).

**Figure S4.**
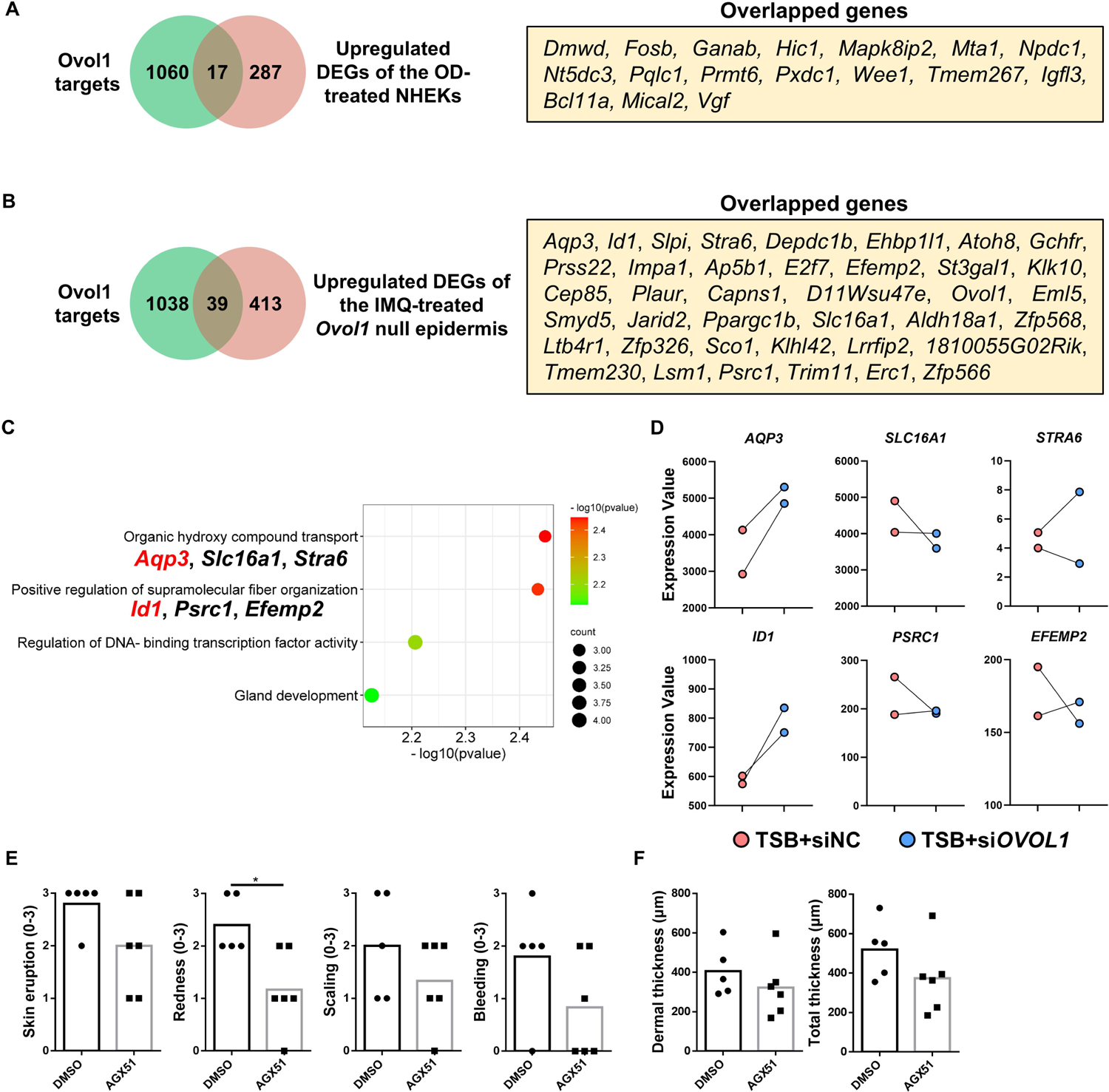
Supporting evidence for *Id1* as a direct and functional target of Ovol1 in epidermal keratinocytes. Related to **Figure 4. (A)** Venn diagram of Ovol1 target genes and genes upregulated in the OD (si*OVOL1* + DMSO)-treated NHEKs compared to ND (siNC + DMSO)-treated NHEKs. **(B)** Venn diagram of Ovol1 target genes and genes upregulated in the IMQ-treated *Ovol1* null epidermis compared to control littermate. For both (A) and (B), gene numbers are indicated in the Venn diagram, and overlapping genes are listed on the left. **(C)** Ingenuity pathways analysis of the 39 overlapping genes from (B). **(D)** RNA-seq results for the indicated genes in TSB-treated NHEKs with or without *OVOL1* depletion. **(E)** Clinical scores of skin eruption, scaling, bleeding and redness of DMSO- or AGX51-treated SSKO mice at day 11. **(F)** Quantification of dermal and total skin thickness. n = 5-6 mice per group for (E-F). * *p* < 0.05. p values were calculated using 2-tailed unpaired Student *t* test., OD: si*OVOL1* + DMSO, ND: siNC + DMSO, TSB: Tryptic Soy Broth.

**Figure S5.**
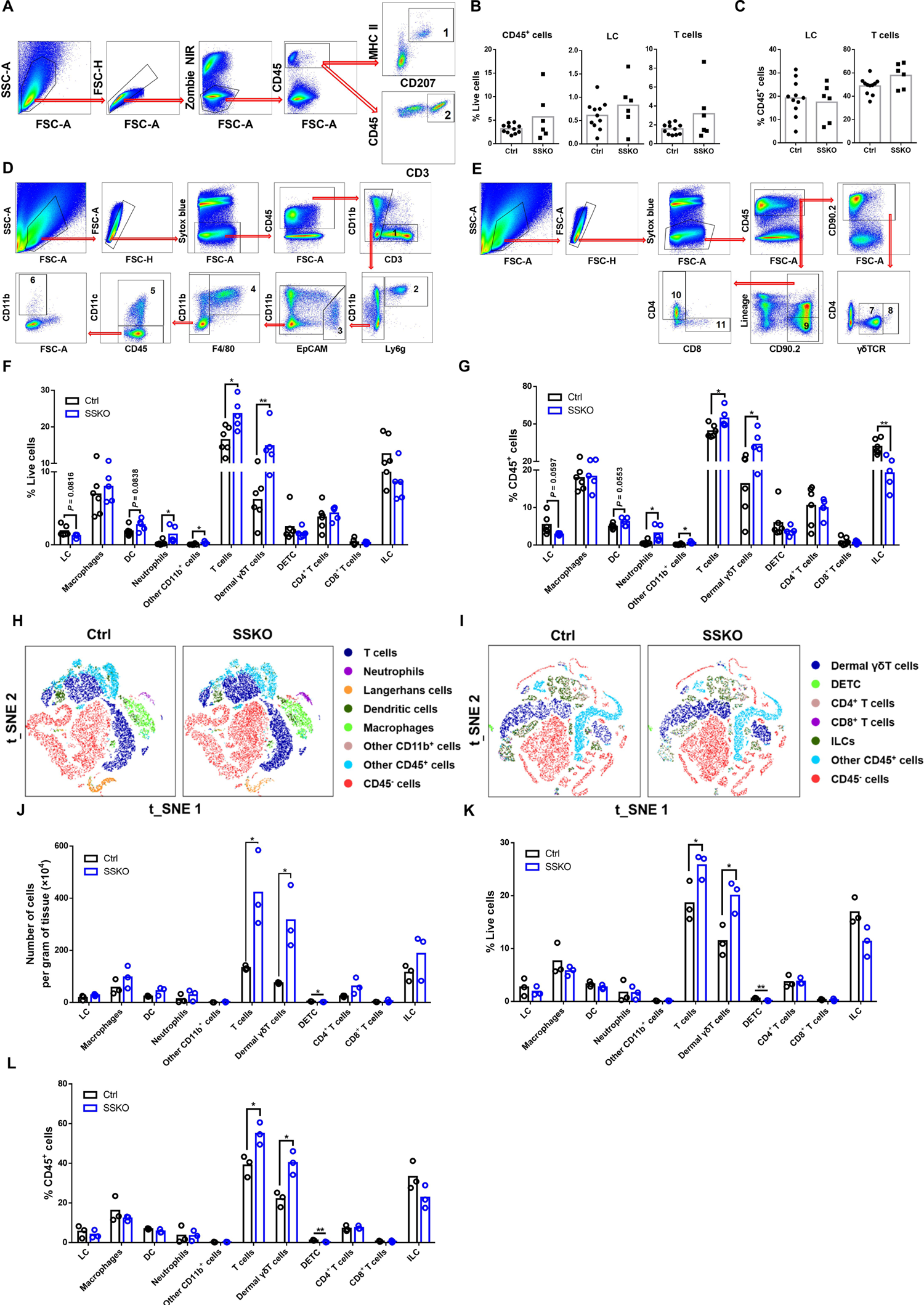
DCs, neutrophils and dermal γδT cells are increased in the skin lesions of HDM/SEB-treated SSKO mice. Related to **Figure 5. (A)** Representative flow cytometry plots showing gating strategy for LCs (1) and T cells (2) within the epidermis. **(B)** Percentages of CD45^+^ cells, LCs, and T cells out of live epidermal cells. **(C)** Percentages of LCs and T cells out of epidermal CD45^+^ cells. For (B-C), n = 11 mice for Ctrl group; n = 6 mice for SSKO group. **(D)** Representative flow cytometry plots showing gating strategy for (1) T cells, (2) neutrophils, (3) LCs, (4) macrophages, (5) DCs, and (6) other CD11b^+^ cells. **(E)** Representative flow cytometry plots showing gating strategy for (7) dermal γδT cells, (8) DETCs, (9) ILCs, (10) CD4^+^ T cells, and (11) CD8^+^ T cells. **(F)** Percentages of different immune cell populations out of total skin cells from mice treated with HDM/SEB at day 11. **(G)** Percentages of different immune cell populations out of skin CD45^+^ cells from mice treated with HDM/SEB at day 11. n = 5-6 mice per group (F, G). **(H-I)** tSNE visualization of different immune cell populations in mice treated with HDM/SEB at day 31. **(J)** Quantification of the indicated immune cell types per gram of skin tissue from mice treated with HDM/SEB at day 31. n = 3 mice per group. **(K-L)** Percentages of different immune cells out of total skin cells (K) or skin CD45^+^ cells (L) from mice treated with HDM/SEB at day 31. n = 3 mice per group. * *p* < 0.05, ** *p* < 0.01. *p* values were calculated using 2-tailed unpaired Student *t* test.

**Figure S6.**
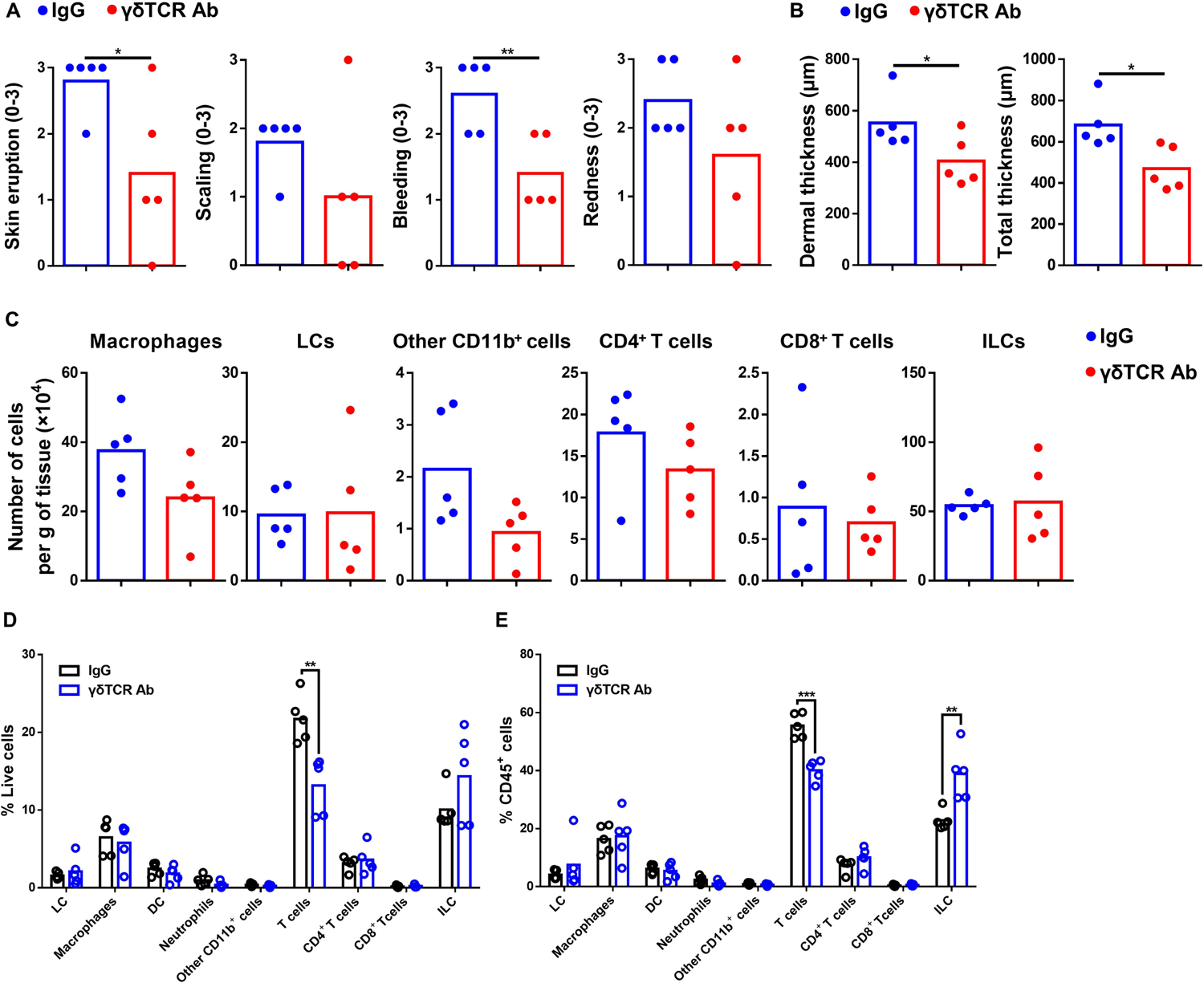
Supporting evidence for the importance of γδT cells in skin inflammation of HDM/SEB-treated SSKO mice. Related to **Figure 6. (A)** Clinical scores of skin eruption, scaling, bleeding and redness in IgG- or γδTCR Ab-treated SSKO mice at day 11. **(B)** Quantification of dermal and total skin thickness in IgG- or γδTCR Ab-treated SSKO mice at day 11. **(C)** Quantification of the indicated immune cell populations per gram of skin tissue at day 11. **(D-E)** Percentages of different immune cells out of total skin cells (D) or skin CD45^+^ cells (E) at day 11. n = 5 mice per group. * *p* < 0.05, ** *p* < 0.01, ***, *p* < 0.001. *p* values were calculated using 2-tailed unpaired Student *t* test.

**Figure S7.**
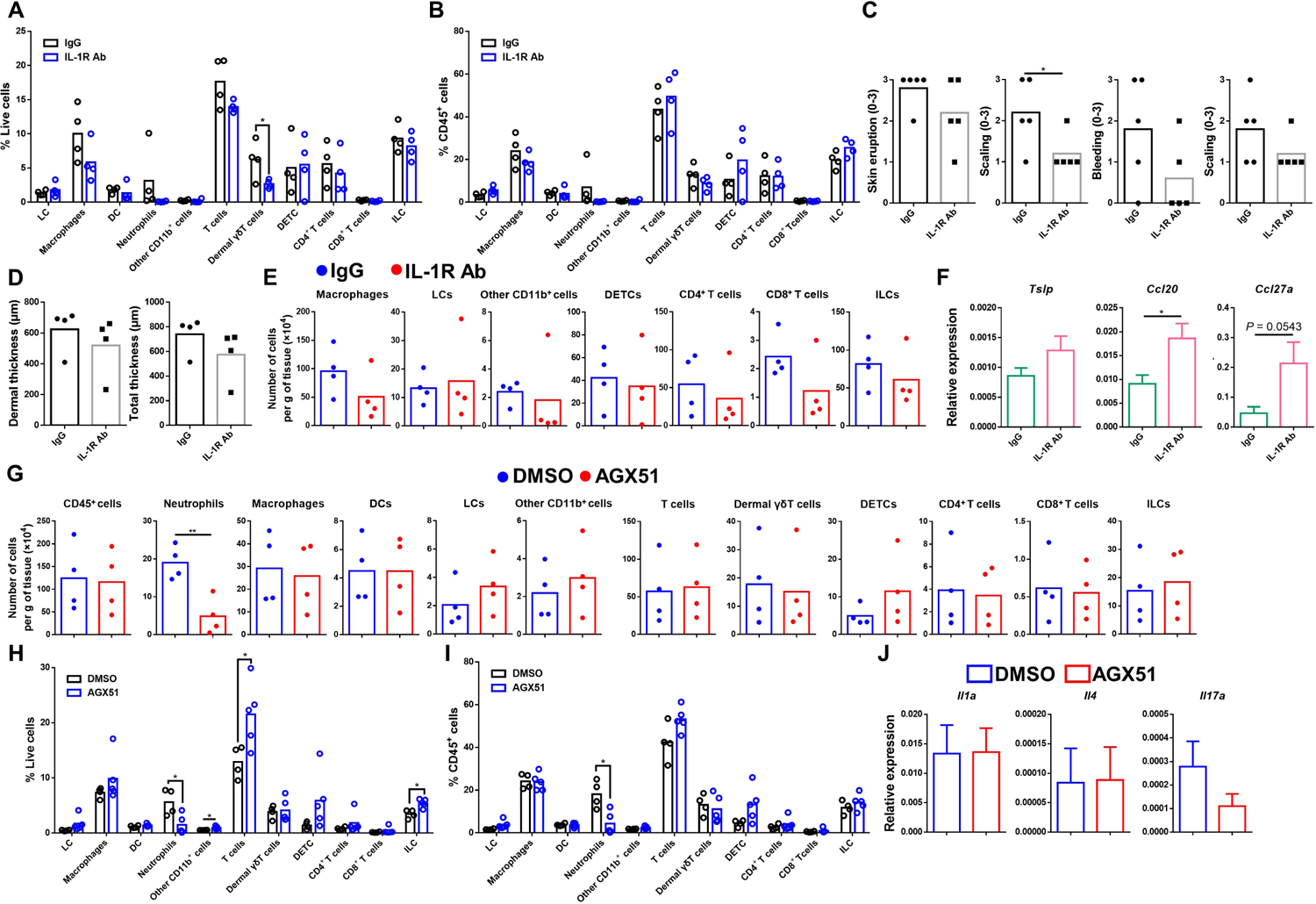
Supporting evidence for the differential roles of IL-1 signaling and Id1 in skin inflammation of HDM/SEB-treated SSKO mice. Related to **Figure 7. (A-B)** Percentages of different immune cell populations out of total skin cells (A) or skin CD45^+^ cells (B) in IgG- or IL-1R Ab-treated SSKO mice at day 11. n = 4 mice per group. **(C)** Clinical scores of skin eruption, scaling, bleeding and redness at day 11. n = 5 mice per group. **(D)** Quantification of dermal and total skin thickness. n = 4 mice per group. **(E)** Quantification of the indicated immune cell types per gram of skin tissue at day 11. n = 4 mice per group. **(F)** RT-qPCR analysis of the indicated genes in whole skin at day 11. n = 5 mice per group. **(G)** Quantification of the indicated immune cell types per gram of skin tissue in DMSO- or AGX51-treated SSKO mice at day 11. n = 4 mice per group. **(H-I)** Percentages of different immune cell populations out of total skin cells (H) or skin CD45^+^ cells (I) at day 11. n = 4-5 mice per group. **(J)** RT-qPCR results of the indicated genes in the whole skin at day 11. Data are mean + SEM. n = 4 mice per group. * *p* < 0.05, ** *p* < 0.01. *p* values were calculated using 2-tailed unpaired Student *t* test.

**Figure S8.**
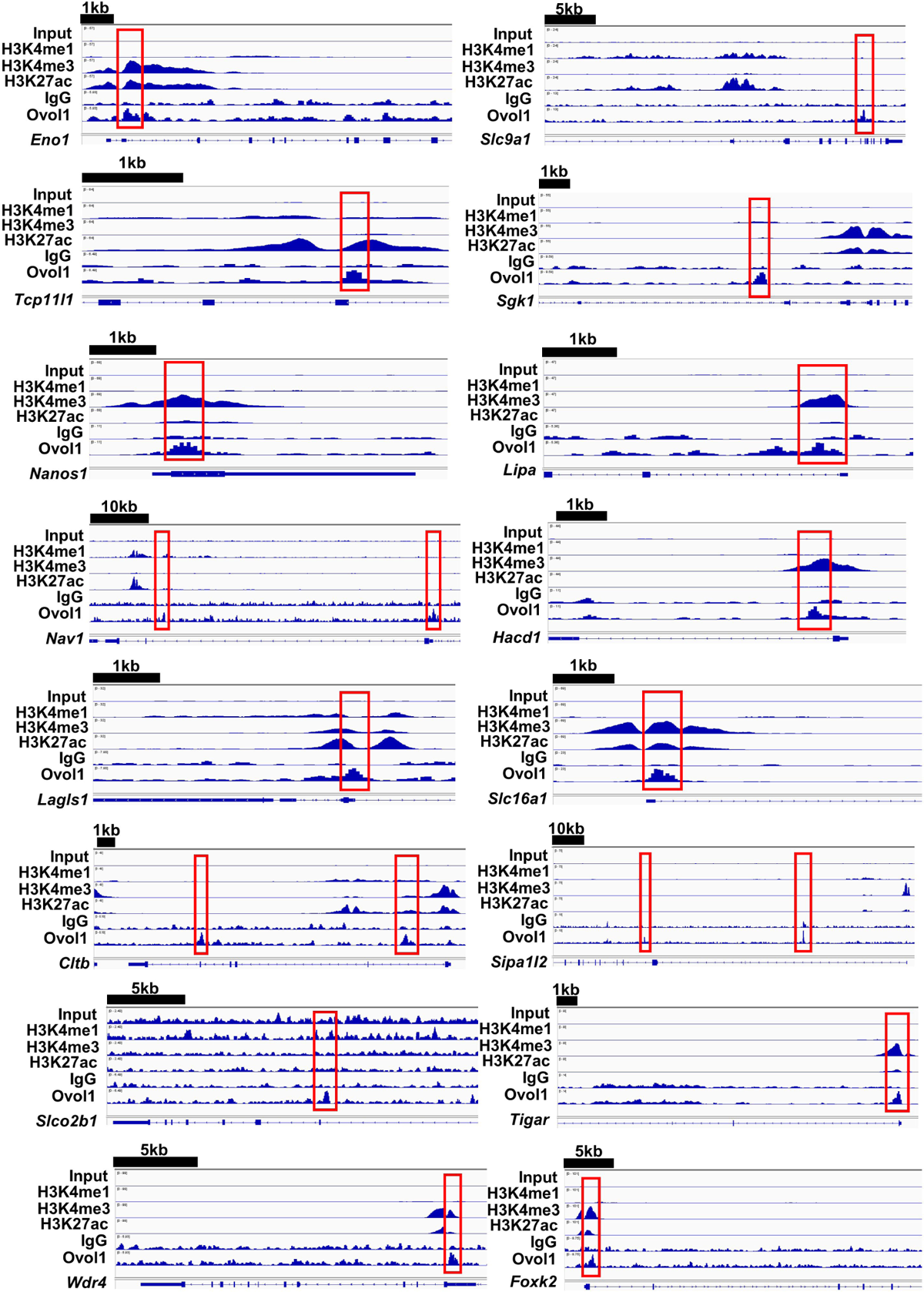
Genome browser track for the indicated ChIP-seq signals across the indicated Ovol1 target gene loci. Related to **Figure 8**. Red boxes highlight the Ovol1-bound regions.

## REFERENCES

1. T. Kobayashi, S. Naik, K. Nagao, Choreographing Immunity in the Skin Epithelial Barrier. Immunity 50, 552–565 (2019).

2. E. Goleva, E. Berdyshev, D. Y. Leung, Epithelial barrier repair and prevention of allergy. J Clin Invest 129, 1463–1474 (2019).

3. S. Stander, Atopic Dermatitis. N Engl J Med 384, 1136–1143 (2021).

4. T. Dainichi, A. Kitoh, A. Otsuka, S. Nakajima, T. Nomura, D. H. Kaplan, K. Kabashima, The epithelial immune microenvironment (EIME) in atopic dermatitis and psoriasis. Nat Immunol 19, 1286–1298 (2018).

5. S. Weidinger, L. A. Beck, T. Bieber, K. Kabashima, A. D. Irvine, Atopic dermatitis. Nat Rev Dis Primers 4, 1 (2018).

6. F. G. Quiroz, V. F. Fiore, J. Levorse, L. Polak, E. Wong, H. A. Pasolli, E. Fuchs, Liquid-liquid phase separation drives skin barrier formation. Science 367, (2020).

7. C. Zhang, M. Chinnappan, C. A. Prestwood, M. Edwards, M. Artami, B. M. Thompson, K. M. Eckert, G. Vale, C. C. Zouboulis, J. G. McDonald, T. A. Harris-Tryon, Interleukins 4 and 13 drive lipid abnormalities in skin cells through regulation of sex steroid hormone synthesis. Proc Natl Acad Sci U S A 118, (2021).

8. Y. Mitamura, S. Nunomura, Y. Nanri, M. Ogawa, T. Yoshihara, M. Masuoka, G. Tsuji, T. Nakahara, A. Hashimoto-Hachiya, S. J. Conway, M. Furue, K. Izuhara, The IL-13/periostin/IL-24 pathway causes epidermal barrier dysfunction in allergic skin inflammation. Allergy 73, 1881–1891 (2018).

9. M. Omori-Miyake, M. Yamashita, Y. Tsunemi, M. Kawashima, J. Yagi, In vitro assessment of IL-4- or IL-13-mediated changes in the structural components of keratinocytes in mice and humans. J Invest Dermatol 134, 1342–1350 (2014).

10. F. Annunziato, C. Romagnani, S. Romagnani, The 3 major types of innate and adaptive cell-mediated effector immunity. J Allergy Clin Immunol 135, 626–635 (2015).

11. L. K. Oetjen, M. R. Mack, J. Feng, T. M. Whelan, H. Niu, C. J. Guo, S. Chen, A. M. Trier, A. Z. Xu, S. V. Tripathi, J. Luo, X. Gao, L. Yang, S. L. Hamilton, P. L. Wang, J. R. Brestoff, M. L. Council, R. Brasington, A. Schaffer, F. Brombacher, C. S. Hsieh, R. W. t. Gereau, M. J. Miller, Z. F. Chen, H. Hu, S. Davidson, Q. Liu, B. S. Kim, Sensory Neurons Co-opt Classical Immune Signaling Pathways to Mediate Chronic Itch. Cell 171, 217–228 e213 (2017).

12. K. Ahn, B. E. Kim, J. Kim, D. Y. Leung, Recent advances in atopic dermatitis. Current opinion in immunology 66, 14–21 (2020).

13. P. M. Brunner, D. Y. M. Leung, E. Guttman-Yassky, Immunologic, microbial, and epithelial interactions in atopic dermatitis. *Annals of allergy, asthma & immunology: official publication of the American College of Allergy, Asthma*, & Immunology 120, 34–41 (2018).

14. P. Y. Ong, Atopic dermatitis: Is innate or adaptive immunity in control? A clinical perspective. Frontiers in immunology 13, 943640 (2022).

15. S. Nakajima, A. Kitoh, G. Egawa, Y. Natsuaki, S. Nakamizo, C. S. Moniaga, A. Otsuka, T. Honda, S. Hanakawa, W. Amano, Y. Iwakura, S. Nakae, M. Kubo, Y. Miyachi, K. Kabashima, IL-17A as an inducer for Th2 immune responses in murine atopic dermatitis models. J Invest Dermatol 134, 2122–2130 (2014).

16. N. Yang, Z. Chen, X. Zhang, Y. Shi, Novel Targeted Biological Agents for the Treatment of Atopic Dermatitis. BioDrugs 35, 401–415 (2021).

17. A. M. Trier, B. S. Kim, Insights into atopic dermatitis pathogenesis lead to newly approved systemic therapies. The British journal of dermatology 188, 698–708 (2023).

18. M. G. Lebwohl, L. Stein Gold, B. Strober, K. A. Papp, A. W. Armstrong, J. Bagel, L. Kircik, B. Ehst, H. C. Hong, J. Soung, J. Fromowitz, S. Guenthner, S. C. Piscitelli, D. S. Rubenstein, P. M. Brown, A. M. Tallman, R. Bissonnette, Phase 3 Trials of Tapinarof Cream for Plaque Psoriasis. N Engl J Med 385, 2219–2229 (2021).

19. A. Uberoi, C. Bartow-McKenney, Q. Zheng, L. Flowers, A. Campbell, S. A. B. Knight, N. Chan, M. Wei, V. Lovins, J. Bugayev, J. Horwinski, C. Bradley, J. Meyer, D. Crumrine, C. H. Sutter, P. Elias, E. Mauldin, T. R. Sutter, E. A. Grice, Commensal microbiota regulates skin barrier function and repair via signaling through the aryl hydrocarbon receptor. Cell Host Microbe 29, 1235–1248 e1238 (2021).

20. K. Haas, H. Weighardt, R. Deenen, K. Köhrer, B. Clausen, S. Zahner, P. Boukamp, W. Bloch, J. Krutmann, C. Esser, Aryl Hydrocarbon Receptor in Keratinocytes Is Essential for Murine Skin Barrier Integrity. J Invest Dermatol 136, 2260–2269 (2016).

21. P. Di Meglio, J. H. Duarte, H. Ahlfors, N. D. Owens, Y. Li, F. Villanova, I. Tosi, K. Hirota, F. O. Nestle, U. Mrowietz, M. J. Gilchrist, B. Stockinger, Activation of the aryl hydrocarbon receptor dampens the severity of inflammatory skin conditions. Immunity 40, 989–1001 (2014).

22. S. H. Smith, C. Jayawickreme, D. J. Rickard, E. Nicodeme, T. Bui, C. Simmons, C. M. Coquery, J. Neil, W. M. Pryor, D. Mayhew, D. K. Rajpal, K. Creech, S. Furst, J. Lee, D. Wu, F. Rastinejad, T. M. Willson, F. Viviani, D. C. Morris, J. T. Moore, J. Cote-Sierra, Tapinarof Is a Natural AhR Agonist that Resolves Skin Inflammation in Mice and Humans. J Invest Dermatol 137, 2110–2119 (2017).

23. E. H. van den Bogaard, J. G. Bergboer, M. Vonk-Bergers, I. M. van Vlijmen-Willems, S. V. Hato, P. G. van der Valk, J. M. Schröder, I. Joosten, P. L. Zeeuwen, J. Schalkwijk, Coal tar induces AHR-dependent skin barrier repair in atopic dermatitis. J Clin Invest 123, 917–927 (2013).

24. A. S. Paller, L. Stein Gold, J. Soung, A. M. Tallman, D. S. Rubenstein, M. Gooderham, Efficacy and patient-reported outcomes from a phase 2b, randomized clinical trial of tapinarof cream for the treatment of adolescents and adults with atopic dermatitis. J Am Acad Dermatol 84, 632–638 (2021).

25. J. Yu, Y. Luo, Z. Zhu, Y. Zhou, L. Sun, J. Gao, J. Sun, G. Wang, X. Yao, W. Li, A tryptophan metabolite of the skin microbiota attenuates inflammation in patients with atopic dermatitis through the aryl hydrocarbon receptor. J Allergy Clin Immunol 143, 2108–2119.e2112 (2019).

26. G. Tsuji, A. Hashimoto-Hachiya, M. Kiyomatsu-Oda, M. Takemura, F. Ohno, T. Ito, S. Morino-Koga, C. Mitoma, T. Nakahara, H. Uchi, M. Furue, Aryl hydrocarbon receptor activation restores filaggrin expression via OVOL1 in atopic dermatitis. Cell Death Dis 8, e2931 (2017).

27. L. Paternoster, M. Standl, C. M. Chen, A. Ramasamy, K. Bonnelykke, L. Duijts, M. A. Ferreira, A. C. Alves, J. P. Thyssen, E. Albrecht, H. Baurecht, B. Feenstra, P. M. Sleiman, P. Hysi, N. M. Warrington, I. Curjuric, R. Myhre, J. A. Curtin, M. M. Groen-Blokhuis, M. Kerkhof, A. Saaf, A. Franke, D. Ellinghaus, R. Folster-Holst, E. Dermitzakis, S. B. Montgomery, H. Prokisch, K. Heim, A. L. Hartikainen, A. Pouta, J. Pekkanen, A. I. Blakemore, J. L. Buxton, M. Kaakinen, D. L. Duffy, P. A. Madden, A. C. Heath, G. W. Montgomery, P. J. Thompson, M. C. Matheson, P. Le Souef, C. Australian Asthma Genetics, B. St Pourcain, G. D. Smith, J. Henderson, J. P. Kemp, N. J. Timpson, P. Deloukas, S. M. Ring, H. E. Wichmann, M. Muller-Nurasyid, N. Novak, N. Klopp, E. Rodriguez, W. McArdle, A. Linneberg, T. Menne, E. A. Nohr, A. Hofman, A. G. Uitterlinden, C. M. van Duijn, F. Rivadeneira, J. C. de Jongste, R. J. van der Valk, M. Wjst, R. Jogi, F. Geller, H. A. Boyd, J. C. Murray, C. Kim, F. Mentch, M. March, M. Mangino, T. D. Spector, V. Bataille, C. E. Pennell, P. G. Holt, P. Sly, C. M. Tiesler, E. Thiering, T. Illig, M. Imboden, W. Nystad, A. Simpson, J. J. Hottenga, D. Postma, G. H. Koppelman, H. A. Smit, C. Soderhall, B. Chawes, E. Kreiner-Moller, H. Bisgaard, E. Melen, D. I. Boomsma, A. Custovic, B. Jacobsson, N. M. Probst-Hensch, L. J. Palmer, D. Glass, H. Hakonarson, M. Melbye, D. L. Jarvis, V. W. Jaddoe, C. Gieger, C. Genetics of Overweight Young Adults, D. P. Strachan, N. G. Martin, M. R. Jarvelin, J. Heinrich, D. M. Evans, S. Weidinger, E. A. Genetics, C. Lifecourse Epidemiology, Meta-analysis of genome-wide association studies identifies three new risk loci for atopic dermatitis. Nat Genet 44, 187–192 (2011).

28. T. Hirota, A. Takahashi, M. Kubo, T. Tsunoda, K. Tomita, M. Sakashita, T. Yamada, S. Fujieda, S. Tanaka, S. Doi, A. Miyatake, T. Enomoto, C. Nishiyama, N. Nakano, K. Maeda, K. Okumura, H. Ogawa, S. Ikeda, E. Noguchi, T. Sakamoto, N. Hizawa, K. Ebe, H. Saeki, T. Sasaki, T. Ebihara, M. Amagai, S. Takeuchi, M. Furue, Y. Nakamura, M. Tamari, Genome-wide association study identifies eight new susceptibility loci for atopic dermatitis in the Japanese population. Nat Genet 44, 1222–1226 (2012).

29. M. Nair, A. Teng, V. Bilanchone, A. Agrawal, B. Li, X. Dai, Ovol1 regulates the growth arrest of embryonic epidermal progenitor cells and represses c-myc transcription. J Cell Biol 173, 253–264 (2006).

30. M. Dragan, P. Sun, Z. Chen, X. Ma, R. Vu, Y. Shi, S. A. Villalta, X. Dai, Epidermis-Intrinsic Transcription Factor Ovol1 Coordinately Regulates Barrier Maintenance and Neutrophil Accumulation in Psoriasis-Like Inflammation. J Invest Dermatol 142, 583–593 e585 (2022).

31. P. Sun, R. Vu, M. Dragan, D. Haensel, G. Gutierrez, Q. Nguyen, E. Greenberg, Z. Chen, J. Wu, S. Atwood, E. Pearlman, Y. Shi, W. Han, K. Kessenbrock, X. Dai, OVOL1 Regulates Psoriasis-Like Skin Inflammation and Epidermal Hyperplasia. J Invest Dermatol 141, 1542–1552 (2021).

32. M. Dragan, Z. Chen, Y. Li, J. Le, P. Sun, D. Haensel, S. Sureshchandra, A. Pham, E. Lu, K. T. Pham, A. Verlande, R. Vu, G. Gutierrez, W. Li, C. Jang, S. Masri, X. Dai, Ovol1/2 loss-induced epidermal defects elicit skin immune activation and alter global metabolism. EMBO reports 24, e56214 (2023).

33. V. Rothhammer, F. J. Quintana, The aryl hydrocarbon receptor: an environmental sensor integrating immune responses in health and disease. Nature reviews. Immunology 19, 184–197 (2019).

34. H. D. Zomer, A. G. Trentin, Skin wound healing in humans and mice: Challenges in translational research. Journal of dermatological science 90, 3–12 (2018).

35. M. D. Lynch, F. M. Watt, Fibroblast heterogeneity: implications for human disease. J Clin Invest 128, 26–35 (2018).

36. T. Hidaka, E. Ogawa, E. H. Kobayashi, T. Suzuki, R. Funayama, T. Nagashima, T. Fujimura, S. Aiba, K. Nakayama, R. Okuyama, M. Yamamoto, The aryl hydrocarbon receptor AhR links atopic dermatitis and air pollution via induction of the neurotrophic factor artemin. Nat Immunol 18, 64–73 (2017).

37. E. H. van den Bogaard, M. A. Podolsky, J. P. Smits, X. Cui, C. John, K. Gowda, D. Desai, S. G. Amin, J. Schalkwijk, G. H. Perdew, A. B. Glick, Genetic and pharmacological analysis identifies a physiological role for the AHR in epidermal differentiation. J Invest Dermatol 135, 1320–1328 (2015).

38. T. Matikainen, G. I. Perez, A. Jurisicova, J. K. Pru, J. J. Schlezinger, H. Y. Ryu, J. Laine, T. Sakai, S. J. Korsmeyer, R. F. Casper, D. H. Sherr, J. L. Tilly, Aromatic hydrocarbon receptor-driven Bax gene expression is required for premature ovarian failure caused by biohazardous environmental chemicals. Nat Genet 28, 355–360 (2001).

39. I. Y. Rojas, B. J. Moyer, C. S. Ringelberg, O. M. Wilkins, D. B. Pooler, D. B. Ness, S. Coker, T. D. Tosteson, L. D. Lewis, M. D. Chamberlin, C. R. Tomlinson, Kynurenine-Induced Aryl Hydrocarbon Receptor Signaling in Mice Causes Body Mass Gain, Liver Steatosis, and Hyperglycemia. *Obesity (Silver Spring*, Md.) 29, 337–349 (2021).

40. H. Lu, W. Cui, C. D. Klaassen, Nrf2 protects against 2,3,7,8-tetrachlorodibenzo-p-dioxin (TCDD)-induced oxidative injury and steatohepatitis. Toxicology and applied pharmacology 256, 122–135 (2011).

41. M. Kiyomatsu-Oda, H. Uchi, S. Morino-Koga, M. Furue, Protective role of 6-formylindolo[3,2-b]carbazole (FICZ), an endogenous ligand for arylhydrocarbon receptor, in chronic mite-induced dermatitis. Journal of dermatological science 90, 284–294 (2018).

42. H. Esaki, D. A. Ewald, B. Ungar, M. Rozenblit, X. Zheng, H. Xu, Y. D. Estrada, X. Peng, H. Mitsui, T. Litman, M. Suárez-Fariñas, J. G. Krueger, E. Guttman-Yassky, Identification of novel immune and barrier genes in atopic dermatitis by means of laser capture microdissection. J Allergy Clin Immunol 135, 153–163 (2015).

43. N. Serhan, L. Basso, R. Sibilano, C. Petitfils, J. Meixiong, C. Bonnart, L. L. Reber, T. Marichal, P. Starkl, N. Cenac, X. Dong, M. Tsai, S. J. Galli, N. Gaudenzio, House dust mites activate nociceptor-mast cell clusters to drive type 2 skin inflammation. Nat Immunol 20, 1435–1443 (2019).

44. G. Reynolds, P. Vegh, J. Fletcher, E. F. M. Poyner, E. Stephenson, I. Goh, R. A. Botting, N. Huang, B. Olabi, A. Dubois, D. Dixon, K. Green, D. Maunder, J. Engelbert, M. Efremova, K. Polanski, L. Jardine, C. Jones, T. Ness, D. Horsfall, J. McGrath, C. Carey, D. M. Popescu, S. Webb, X. N. Wang, B. Sayer, J. E. Park, V. A. Negri, D. Belokhvostova, M. D. Lynch, D. McDonald, A. Filby, T. Hagai, K. B. Meyer, A. Husain, J. Coxhead, R. Vento-Tormo, S. Behjati, S. Lisgo, A. C. Villani, J. Bacardit, P. H. Jones, E. A. O’Toole, G. S. Ogg, N. Rajan, N. J. Reynolds, S. A. Teichmann, F. M. Watt, M. Haniffa, Developmental cell programs are co-opted in inflammatory skin disease. Science 371, (2021).

45. V. Moosbrugger-Martinz, M. Schmuth, S. Dubrac, A Mouse Model for Atopic Dermatitis Using Topical Application of Vitamin D3 or of Its Analog MC903. Methods in molecular biology (Clifton, N.J.) 1559, 91–106 (2017).

46. W. Mao, A. C. Salzberg, M. Uchigashima, Y. Hasegawa, H. Hock, M. Watanabe, S. Akbarian, Y. I. Kawasawa, K. Futai, Activity-Induced Regulation of Synaptic Strength through the Chromatin Reader L3mbtl1. Cell Rep 23, 3209–3222 (2018).

47. H. Alam, L. Sehgal, S. T. Kundu, S. N. Dalal, M. M. Vaidya, Novel function of keratins 5 and 14 in proliferation and differentiation of stratified epithelial cells. Molecular biology of the cell 22, 4068–4078 (2011).

48. M. Hara-Chikuma, H. Satooka, S. Watanabe, T. Honda, Y. Miyachi, T. Watanabe, A. S. Verkman, Aquaporin-3-mediated hydrogen peroxide transport is required for NF-kappaB signalling in keratinocytes and development of psoriasis. Nat Commun 6, 7454 (2015).

49. Y. Hamajima, M. Komori, D. A. Preciado, D. I. Choo, K. Moribe, S. Murakami, F. G. Ondrey, J. Lin, The role of inhibitor of DNA-binding (Id1) in hyperproliferation of keratinocytes: the pathological basis for middle ear cholesteatoma from chronic otitis media. Cell Prolif 43, 457–463 (2010).

50. S. J. Renaud, D. Chakraborty, C. W. Mason, M. A. Rumi, J. L. Vivian, M. J. Soares, OVO-like 1 regulates progenitor cell fate in human trophoblast development. Proc Natl Acad Sci U S A 112, E6175–6184 (2015).

51. C. G. Kantzer, W. Yang, D. Grommisch, K. Vikhe Patil, K. H. Mak, V. Shirokova, M. Genander, ID1 and CEBPA coordinate epidermal progenitor cell differentiation. *Development (Cambridge*, England) 149, (2022).

52. P. M. Wojnarowicz, E. S. R. Lima, M. Ohnaka, S. B. Lee, Y. Chin, A. Kulukian, S. H. Chang, B. Desai, M. Garcia Escolano, R. Shah, M. Garcia-Cao, S. Xu, R. Kadam, Y. Goldgur, M. A. Miller, O. Ouerfelli, G. Yang, T. Arakawa, S. K. Albanese, W. A. Garland, G. Stoller, J. Chaudhary, L. Norton, R. K. Soni, J. Philip, R. C. Hendrickson, A. Iavarone, A. J. Dannenberg, J. D. Chodera, N. Pavletich, A. Lasorella, P. A. Campochiaro, R. Benezra, A Small-Molecule Pan-Id Antagonist Inhibits Pathologic Ocular Neovascularization. Cell Rep 29, 62–75 e67 (2019).

53. H. C. Hawerkamp, C. M. R. Fahy, P. G. Fallon, C. Schwartz, Break on through: The role of innate immunity and barrier defence in atopic dermatitis and psoriasis. Skin health and disease 2, e99 (2022).

54. P. L. Tong, B. Roediger, N. Kolesnikoff, M. Biro, S. S. Tay, R. Jain, L. E. Shaw, M. A. Grimbaldeston, W. Weninger, The skin immune atlas: three-dimensional analysis of cutaneous leukocyte subsets by multiphoton microscopy. J Invest Dermatol 135, 84–93 (2015).

55. N. Sumaria, B. Roediger, L. G. Ng, J. Qin, R. Pinto, L. L. Cavanagh, E. Shklovskaya, B. Fazekas de St Groth, J. A. Triccas, W. Weninger, Cutaneous immunosurveillance by self-renewing dermal gammadelta T cells. J Exp Med 208, 505–518 (2011).

56. P. H. Papotto, J. C. Ribot, B. Silva-Santos, IL-17(+) γδ T cells as kick-starters of inflammation. Nat Immunol 18, 604–611 (2017).

57. Y. Cai, X. Shen, C. Ding, C. Qi, K. Li, X. Li, V. R. Jala, H. G. Zhang, T. Wang, J. Zheng, J. Yan, Pivotal role of dermal IL-17-producing gammadelta T cells in skin inflammation. Immunity 35, 596–610 (2011).

58. C. Koenecke, V. Chennupati, S. Schmitz, B. Malissen, R. Forster, I. Prinz, In vivo application of mAb directed against the gammadelta TCR does not deplete but generates “invisible” gammadelta T cells. Eur J Immunol 39, 372–379 (2009).

59. Y. Han, J. Mora, A. Huard, P. da Silva, S. Wiechmann, M. Putyrski, C. Schuster, E. Elwakeel, G. Lang, A. Scholz, T. Scholz, T. Schmid, N. de Bruin, P. Billuart, C. Sala, H. Burkhardt, M. J. Parnham, A. Ernst, B. Brune, A. Weigert, IL-38 Ameliorates Skin Inflammation and Limits IL-17 Production from gammadelta T Cells. Cell Rep 27, 835–846 e835 (2019).

60. Y. S. Chen, I. B. Chen, G. Pham, T. Y. Shao, H. Bangar, S. S. Way, D. B. Haslam, IL-17-producing gammadelta T cells protect against Clostridium difficile infection. J Clin Invest 130, 2377–2390 (2020).

61. R. L. O’Brien, W. K. Born, Dermal γδ T cells--What have we learned? Cellular immunology 296, 62–69 (2015).

62. K. A. Engebretsen, J. D. Johansen, S. Kezic, A. Linneberg, J. P. Thyssen, The effect of environmental humidity and temperature on skin barrier function and dermatitis. J Eur Acad Dermatol Venereol 30, 223–249 (2016).

63. L. C. Wood, S. M. Jackson, P. M. Elias, C. Grunfeld, K. R. Feingold, Cutaneous barrier perturbation stimulates cytokine production in the epidermis of mice. J Clin Invest 90, 482–487 (1992).

64. Y. Cai, F. Xue, C. Quan, M. Qu, N. Liu, Y. Zhang, C. Fleming, X. Hu, H. G. Zhang, R. Weichselbaum, Y. X. Fu, D. Tieri, E. C. Rouchka, J. Zheng, J. Yan, A Critical Role of the IL-1beta-IL-1R Signaling Pathway in Skin Inflammation and Psoriasis Pathogenesis. J Invest Dermatol 139, 146–156 (2019).

65. Y. Cai, F. Xue, C. Fleming, J. Yang, C. Ding, Y. Ma, M. Liu, H. G. Zhang, J. Zheng, N. Xiong, J. Yan, Differential developmental requirement and peripheral regulation for dermal Vgamma4 and Vgamma6T17 cells in health and inflammation. Nat Commun 5, 3986 (2014).

66. Y. Z. Liu, M. Y. Xu, X. Y. Dai, L. Yan, L. Li, R. Z. Zhu, L. J. Ren, J. Q. Zhang, X. F. Zhang, J. F. Li, Y. J. Tian, W. J. Shi, Y. Q. Liu, C. L. Jiang, J. B. Zhu, J. K. Chen, Pyruvate Kinase M2 Mediates Glycolysis Contributes to Psoriasis by Promoting Keratinocyte Proliferation. Frontiers in pharmacology 12, 765790 (2021).

67. T. Tohgasaki, N. Ozawa, T. Yoshino, S. Ishiwatari, S. Matsukuma, S. Yanagi, H. Fukuda, Enolase-1 expression in the stratum corneum is elevated with parakeratosis of atopic dermatitis and disrupts the cellular tight junction barrier in keratinocytes. International journal of cosmetic science 40, 178–186 (2018).

68. M. J. Behne, J. W. Meyer, K. M. Hanson, N. P. Barry, S. Murata, D. Crumrine, R. W. Clegg, E. Gratton, W. M. Holleran, P. M. Elias, T. M. Mauro, NHE1 regulates the stratum corneum permeability barrier homeostasis. Microenvironment acidification assessed with fluorescence lifetime imaging. The Journal of biological chemistry 277, 47399–47406 (2002).

69. V. Sukonina, H. Ma, W. Zhang, S. Bartesaghi, S. Subhash, M. Heglind, H. Foyn, M. J. Betz, D. Nilsson, M. E. Lidell, J. Naumann, S. Haufs-Brusberg, H. Palmgren, T. Mondal, M. Beg, M. P. Jedrychowski, K. Taskén, A. Pfeifer, X. R. Peng, C. Kanduri, S. Enerbäck, FOXK1 and FOXK2 regulate aerobic glycolysis. Nature 566, 279–283 (2019).

70. M. Korbelius, K. B. Kuentzel, I. Bradić, N. Vujić, D. Kratky, Recent insights into lysosomal acid lipase deficiency. Trends in molecular medicine 29, 425–438 (2023).

71. A. Flores, J. Schell, A. S. Krall, D. Jelinek, M. Miranda, M. Grigorian, D. Braas, A. C. White, J. L. Zhou, N. A. Graham, T. Graeber, P. Seth, D. Evseenko, H. A. Coller, J. Rutter, H. R. Christofk, W. E. Lowry, Lactate dehydrogenase activity drives hair follicle stem cell activation. Nature cell biology 19, 1017–1026 (2017).

72. E. C. Cheung, R. L. Ludwig, K. H. Vousden, Mitochondrial localization of TIGAR under hypoxia stimulates HK2 and lowers ROS and cell death. Proc Natl Acad Sci U S A 109, 20491–20496 (2012).

73. H. A. Sikder, M. K. Devlin, S. Dunlap, B. Ryu, R. M. Alani, Id proteins in cell growth and tumorigenesis. Cancer cell 3, 525–530 (2003).

74. K. Guilloteau, I. Paris, N. Pedretti, K. Boniface, F. Juchaux, V. Huguier, G. Guillet, F. X. Bernard, J. C. Lecron, F. Morel, Skin Inflammation Induced by the Synergistic Action of IL-17A, IL-22, Oncostatin M, IL-1{alpha}, and TNF-{alpha} Recapitulates Some Features of Psoriasis. Journal of immunology (Baltimore, Md.: 1950) 184, 5263–5270 (2010).

75. N. Fernandez-Gallego, F. Sanchez-Madrid, D. Cibrian, Role of AHR Ligands in Skin Homeostasis and Cutaneous Inflammation. Cells 10, (2021).

76. M. Kyoreva, Y. Li, M. Hoosenally, J. Hardman-Smart, K. Morrison, I. Tosi, M. Tolaini, G. Barinaga, B. Stockinger, U. Mrowietz, F. O. Nestle, C. H. Smith, J. N. Barker, P. Di Meglio, CYP1A1 Enzymatic Activity Influences Skin Inflammation Via Regulation of the AHR Pathway. J Invest Dermatol 141, 1553–1563.e1553 (2021).

77. C. R. Geest, M. Buitenhuis, E. Vellenga, P. J. Coffer, Ectopic expression of C/EBPalpha and ID1 is sufficient to restore defective neutrophil development in low-risk myelodysplasia. Haematologica 94, 1075–1084 (2009).

78. R. Castillo-Gonzalez, D. Cibrian, F. Sanchez-Madrid, Dissecting the complexity of gammadelta T-cell subsets in skin homeostasis, inflammation, and malignancy. J Allergy Clin Immunol 147, 2030–2042 (2021).

79. J. Yoon, J. M. Leyva-Castillo, G. Wang, C. Galand, M. K. Oyoshi, L. Kumar, S. Hoff, R. He, A. Chervonsky, J. J. Oppenheim, V. K. Kuchroo, M. R. van den Brink, W. Malefyt Rde, P. A. Tessier, R. Fuhlbrigge, P. Rosenstiel, C. Terhorst, G. Murphy, R. S. Geha, IL-23 induced in keratinocytes by endogenous TLR4 ligands polarizes dendritic cells to drive IL-22 responses to skin immunization. J Exp Med 213, 2147–2166 (2016).

80. N. Malhotra, J. Yoon, J. M. Leyva-Castillo, C. Galand, N. Archer, L. S. Miller, R. S. Geha, IL-22 derived from gammadelta T cells restricts Staphylococcus aureus infection of mechanically injured skin. J Allergy Clin Immunol 138, 1098–1107 e1093 (2016).

81. N. A. Spidale, N. Malhotra, M. Frascoli, K. Sylvia, B. Miu, C. Freeman, B. D. Stadinski, E. Huseby, J. Kang, Neonatal-derived IL-17 producing dermal gammadelta T cells are required to prevent spontaneous atopic dermatitis. Elife 9, (2020).

82. S. P. Saunders, A. Floudas, T. Moran, C. M. Byrne, M. D. Rooney, C. M. R. Fahy, J. A. Geoghegan, Y. Iwakura, P. G. Fallon, C. Schwartz, Dysregulated skin barrier function in Tmem79 mutant mice promotes IL-17A-dependent spontaneous skin and lung inflammation. Allergy 75, 3216–3227 (2020).

83. D. Tian, Y. Lai, The Relapse of Psoriasis: Mechanisms and Mysteries. JID Innov 2, 100116 (2022).

84. M. Le Borgne, N. Etchart, A. Goubier, S. A. Lira, J. C. Sirard, N. van Rooijen, C. Caux, S. Ait-Yahia, A. Vicari, D. Kaiserlian, B. Dubois, Dendritic cells rapidly recruited into epithelial tissues via CCR6/CCL20 are responsible for CD8+ T cell crosspriming in vivo. Immunity 24, 191–201 (2006).

85. T. R. Matos, J. T. O’Malley, E. L. Lowry, D. Hamm, I. R. Kirsch, H. S. Robins, T. S. Kupper, J. G. Krueger, R. A. Clark, Clinically resolved psoriatic lesions contain psoriasis-specific IL-17-producing alphabeta T cell clones. J Clin Invest 127, 4031–4041 (2017).

86. C. Cairo, E. Arabito, F. Landi, A. Casati, E. Brunetti, G. Mancino, E. Galli, Analysis of circulating gammadelta T cells in children affected by IgE-associated and non-IgE-associated allergic atopic eczema/dermatitis syndrome. Clin Exp Immunol 141, 116–121 (2005).

87. Z. Yu, Y. Gong, L. Cui, Y. Hu, Q. Zhou, Z. Chen, Y. Yu, Y. Chen, P. Xu, X. Zhang, C. Guo, Y. Shi, High-throughput transcriptome and pathogenesis analysis of clinical psoriasis. Journal of dermatological science 98, 109–118 (2020).

88. Y. Zhou, B. Zhou, L. Pache, M. Chang, A. H. Khodabakhshi, O. Tanaseichuk, C. Benner, S. K. Chanda, Metascape provides a biologist-oriented resource for the analysis of systems-level datasets. Nat Commun 10, 1523 (2019).

89. H. Esaki, D. A. Ewald, B. Ungar, M. Rozenblit, X. Zheng, H. Xu, Y. D. Estrada, X. Peng, H. Mitsui, T. Litman, M. Suarez-Farinas, J. G. Krueger, E. Guttman-Yassky, Identification of novel immune and barrier genes in atopic dermatitis by means of laser capture microdissection. J Allergy Clin Immunol 135, 153–163 (2015).

90. D. Haensel, S. Jin, P. Sun, R. Cinco, M. Dragan, Q. Nguyen, Z. Cang, Y. Gong, R. Vu, A. L. MacLean, K. Kessenbrock, E. Gratton, Q. Nie, X. Dai, Defining Epidermal Basal Cell States during Skin Homeostasis and Wound Healing Using Single-Cell Transcriptomics. Cell Rep 30, 3932–3947 e3936 (2020).

91. R. Dai, W. Xu, W. Chen, L. Cui, L. Li, J. Zhou, X. Jin, Y. Wang, L. Wang, Y. Sun, Epigenetic modification of Kiss1 gene expression in the AVPV is essential for female reproductive aging. Bioscience trends 16, 346–358 (2022).

92. Z. Yu, Q. Yu, H. Xu, X. Dai, Y. Yu, L. Cui, Y. Chen, J. Gu, X. Zhang, C. Guo, Y. Shi, IL-17A Promotes Psoriasis-Associated Keratinocyte Proliferation through ACT1-Dependent Activation of YAP-AREG Axis. J Invest Dermatol 142, 2343–2352 (2022).

93. Y. Zhang, T. Liu, C. A. Meyer, J. Eeckhoute, D. S. Johnson, B. E. Bernstein, C. Nusbaum, R. M. Myers, M. Brown, W. Li, X. S. Liu, Model-based analysis of ChIP-Seq (MACS). Genome biology 9, R137 (2008).

94. S. H. Duttke, M. W. Chang, S. Heinz, C. Benner, Identification and dynamic quantification of regulatory elements using total RNA. Genome research 29, 1836–1846 (2019).

